# Chromosomal mutational signatures of DNA damaging agents at single cell resolution

**DOI:** 10.64898/2026.08.30.748106

**Authors:** Mirela Andronescu, Alexander Adrian-Hamazaki, Damian Yap, William Daniels, Armaghan Sarvar, Elena Zaikova, Benjamin Furman, Vinci Au, Amelia Richter, Michael Van Vliet, Caroline Baril, Beixi Wang, Sean Beatty, Farhia Kabeer, Ciara O’Flanagan, Hoa Tran, Viktoriia Cherkasova, BaRun Kim, Teresa Ruiz De Algara, Sophia Means, Nat Westereng, Jay Tan, Jessica Chan, Johnson Zhong, Robert Reinert, Joseph Micla, Benoit Prevost-Potvin, Bita Mojtahedzadeh, Hong Xu, Richard Moore, Andy Mungall, Daniel Lai, Samuel Aparicio

## Abstract

The chromosomal-scale mutational spectrum of small molecules that interact with DNA has been hard to study at scale, as mutational events are distributed in location and occur in parallel in different cells. Here, we present a framework that pairs phylogenetic ancestry reconstruction with mutational signature decomposition to characterise recent, cell-private copy number alteration (CNA) mutational patterns at single-cell resolution. We used this framework to characterise the cell-wise mutational spectrum of contemporaneous CNAs generated by double-strand-break-inducing chemotherapeutic drugs. We demonstrate that platinum salts, G-quadruplex stabilizers and topoisomerase II inhibitors, although mechanistically distinct, converge on a mutational signature dominated by telomere-bounded copy-number gains and losses. This signature is observed in different genetic backgrounds and *in vivo* in drug-treated patient-derived xenografts. We also observe a high rate of endogenous telomere-bounded mutational foreground in *BRCA1* deficient cells. We show that the single cell genome derived signature exposures are drug dose-dependent, and use this to identify the decay of mutational load after drug withdrawal. We observe that both cisplatin and a G4 binder molecule (CX5461) exhibit foreground mutational signature persistence for at least 3 weeks after drug withdrawal, suggesting that residual effects of exposure may last longer than anticipated. Finally, extending the framework to serially drug-treated patient-derived xenograft (PDX) models, we show that telomere-bounded CNA signature exposure is associated with tumoural response to drug, consistent with loss of mutational activity on the genome after acquired resistance emerges. Together, our results show that our framework applied on scWGS identifies contemporaneous chromosomal mutation patterns induced by small molecules in human tissues.

## Introduction

A challenge in decoding the mutational activity of small molecules acting on DNA is that the genomically distributed and parallel nature of mutation obscures observation of mutational activity. Approaches to studying single base or short indel mutations have centered around error-corrected duplex sequencing^1–3^, or cloning and amplification of single cells^4–7^. Duplex sequencing does not effectively capture large scale mutations that occur over megabase scales or the whole genome^3^. Cloning forces cell division and may mask negatively selected mutations^7^. We have previously shown that scaled transposition-based single-cell genome sequencing can capture cell-to-cell private mutations arising from erroneous DNA repair and chromosomal instability mechanisms during replication^8–11^. Here we use DLP+ scWGS to capture private mutations induced by chemicals and identify a common large-scale mutational signature induced by cytotoxic agents used in cancer treatment.

Cytotoxic chemotherapy is a cornerstone of systemic cancer treatment. Different classes of chemotherapy agents induce DNA damage via distinct mechanisms. Platinum-based agents, such as cisplatin^12^, induce intra- and interstrand DNA crosslinks, while CX5461^13^ and preclinical G4 ligand pyridostatin^14^ (PDS) act via stabilising G-quadruplex (G4) DNA structures. Etoposide^15^ and voreloxin^16^ (developed clinically as vosaroxin) are topoisomerase II poisons that stabilise the enzyme-DNA complexes. Due to the distinct double stranded break (DSB)-initiating mechanisms and the various recognition and repair pathways, one would expect these drugs to produce distinct genomic footprints. However, these drug classes converge phenotypically on replication stress^12,17–19^, anaphase bridges and micro-nucleation^20–23^, and chromosomal instability^12,17–23^ raising the possibility that they may share downstream genomic consequences.

The genomic footprints of drug therapy-induced DNA damage have been extensively characterised by bulk single- and double-base substitution (SBS, DBS) mutational signatures^24,25^, including the characteristic signature following cisplatin treatment^26,27^. Furthermore, bulk copy number aberration (CNA) signatures have been shown to be associated with homologous recombination deficiency (HRD) and other genome instability phenotypes^28,29^. However, bulk copy-number profiles are composites of many cells, and are often dominated by recurrent and clonally inherited events, obscuring more recent alterations that occur in single cells. Previous work has developed the concept of mutational foreground: mutations (in our case CNAs) that are private across cells and thereby do not reflect shared clonal history^10^. Existing mutational signature frameworks have not been designed to operate on the foreground layer; hence it remains unclear if different DNA-damaging drug classes retain distinct DNA lesion-specific patterns, or share foreground copy-number patterns based on their shared chromosomal-scale outcomes. It is also not known how such foreground patterns may vary with drug dose, genetic background, treatment withdrawal and tumour sensitivity.

Here we employ phylogenetic reconstruction of cellular ancestry to infer the most recent common ancestor (MRCA) copy-number (CN) state and derive the mutational foreground of each cell by calculating the relative differences from that ancestral state. We then classify foreground copy-number events by segment size, chromosome topology and whether it is a gain or loss. This is then used by the hierarchical Dirichlet process (HDP) to learn recurrent signatures and quantify exposure in individual cells. Using isogenic immortalised and transformed cell lines, we show that treatment by different classes of DNA-damaging agents converges on the telomere-bounded foreground signature; whereas non DNA-damaging drugs do not. The composition and magnitude of this signature is influenced by cellular genetic context, is drug-dose dependent, and its exposure in cells remains detectable for weeks after drug withdrawal. In serially treated patient-derived xenografts (PDX) lines, we observe that the foreground signature exposure changes with tumour response across passages. Taken together, these results demonstrate that CNA foreground signatures are a quantitative genomic readout of treatment-associated chromosomal damage and its dynamics.

### A single-cell phylogenetic framework resolves foreground copy-number changes into mutational signatures

To define the patterns of chromosomal scale mutations induced by DNA damaging small molecules, we took advantage of our previously described approach for transposition-based single-cell whole genome sequencing (scWGS DLP+^9^, Figure 1A). Although transposition based scWGS is shallow in genome coverage, it is non-amplification based and thus produces a more accurate DNA copy number representation in single cells^9^. We treated 184-hTERT-L9 *TP53*-deficient hTERT-immortalized mammary epithelial cells^10^ (*TP53*^-/-^ hTERT) with DNA damaging small molecules that induce DSBs by interacting with DNA via different mechanisms. We first tested cisplatin and CX5461, choosing drug concentrations with detectable effects on cell survival (IC30) but below maximal cell killing (Supplementary Figure 1, Supplementary Table 1). We sequenced post-treated cells with DLP+ scWGS and noted that drug-treated cells tended to have more events compared to untreated immortalized line (Figure 2A,B). Formal investigations into CNAs showed that the total cell-wise mutational event rates (Figure 2C, Extended Data Figure 1) showed minor increases in CX5461-treated samples (generalized linear mixed model (GLMM) fold change = 1.33, *p-adj* = 0.06, Methods), and were significantly higher in cisplatin-treated samples (fold change = 1.74, *p-adj* = 2.15 × 10⁻^4^), with the latter being driven by a rise in whole genome doubled (WGD) cells^30^ (14% → 29%, Methods).

**Figure 1.**
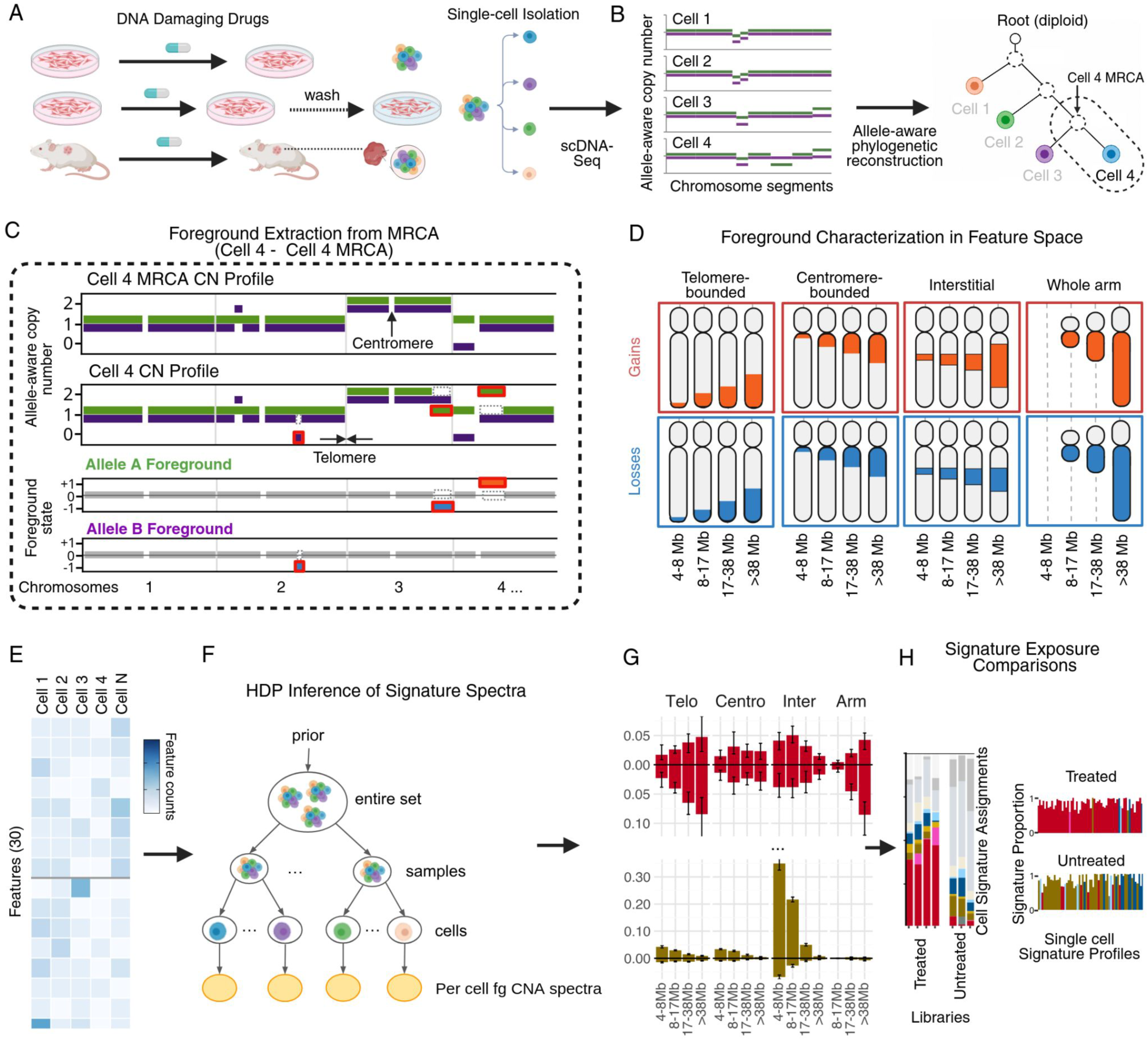
Single-cell analytical framework for resolving cell-specific copy number mutational signatures. **A, Experimental workflow**. *In vitro* cell lines and *in vivo* patient-derived xenograft (PDX) models are subjected to DNA-damaging drug treatments (such as cisplatin, CX5461, etoposide, voreloxin or control drugs) or a given subsequent drug holiday. Single cells are harvested, sorted, and processed via DLP+ single-cell whole-genome sequencing. **B**, **Allele-aware phylogenetic reconstruction.** Allele-specific copy-number (CN) profiles are generated for individual cells, enabling the construction of a CN phylogeny and the inference of each cell’s most recent common ancestor (MRCA) via MEDICC2. **C**, **Mutational foreground extraction.** To separate private non-clonal damage from inherited history, the CN of the cells (terminal leaf of the inferred phylogenetic tree) are subtracted (with allele-awareness) by their inferred MRCA CN state, yielding foreground CN gains (orange) and losses (blue). **D, Foreground feature characterization.** Foreground events larger than 4 Mb are categorized into a multi-dimensional feature space matrix comprising chromosomal topology (telomere-bounded, centromere-bounded, interstitial, whole arm), copy number gains or losses, and discrete size bins, yielding a feature counts matrix with 30 rows per cell **(E)**. **F,G,H**, **Mutational signature analysis.** A hierarchical Dirichlet process (HDP) decomposes the characterized foreground events across thousands of single cells to computationally extract the underlying structural mutational signatures, distinguishing treated from untreated cell populations.

**Figure 2.**
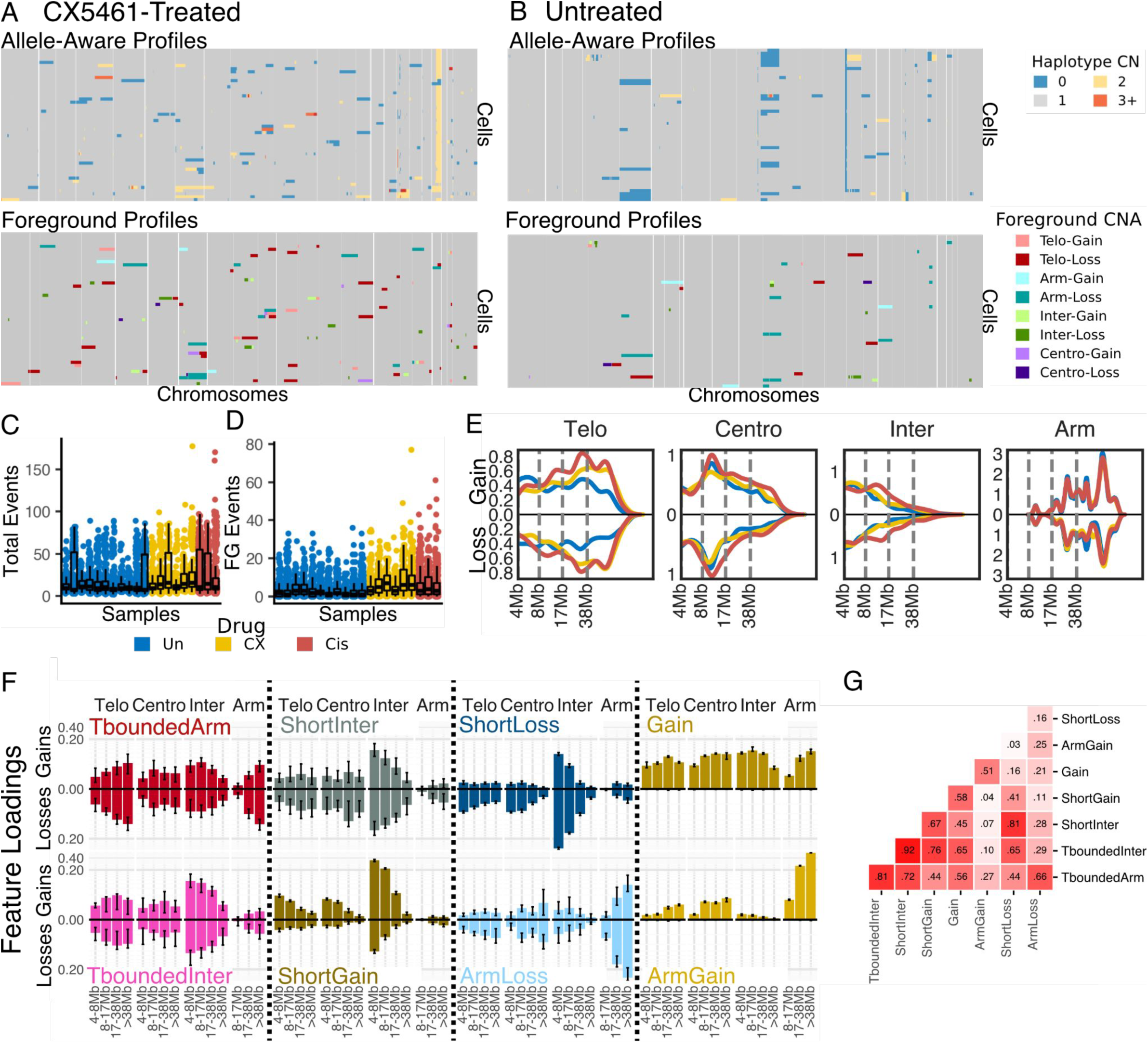
Foreground signature extraction. **A,** Example of absolute allele-aware cell CN profiles (top) for allele A of CX5461-treated *TP53^-/-^*hTERT cells. Example of each cell’s respective foreground CN profiles (bottom), where coloring indicates each CNA’s topology and gain/loss status. **B**, Examples for untreated *TP53^-/-^* hTERTs. **C,** the number of total CNA events, and the number of foreground events **(D)** per cell in untreated, CX5461-treated and cisplatin-treated *TP53^-/-^* hTERT cells. **E**, Densities of foreground events in cells in D as categorized by our feature space. In brief, foreground events are categorized by topology (telomeric-bounded, centromeric-bounded, interstitial, whole arm), size (x axis), or gain (above x axis) or loss (under x axis). **F**, Signature loadings on our 30 features via HDP de-novo decomposition trained on our entire data set. Signatures are named for their dominating characteristics. Error bars represent 95% credible intervals. **G,** Cosine similarities of signature loadings across the features.

As small molecules, or active mutational processes, induce mutations that are initially private to single cells, any method to study them must separate the cell-wise differences (mutational “foreground”) from “background” clone-level events. We thus built single-cell, allele-aware phylogenies, and extracted the events on each allele that were distinct from each cell’s inferred most recent common ancestor (MRCA, Figure 1B,C, Figure 2A,B, Methods). We defined the extracted cell-wise events as the cellular foreground^10^, an assumed mixture of drug and cell-line instability-induced mutational events that are private to individual cells. We focused on events larger than 4 Mb, as these encode the maximum dynamic range across experimental contrasts. We confirm subsequently that inclusion of smaller events does not alter the signature patterns, although it may obscure weaker exposures (Supplementary Figure 2). We investigated foreground event rates (Methods) across drug treatments, finding an increased burden in both CX5461-treated samples (GLMM fold change = 2.37, p-adj = 2.26 x 10⁻^7^) and cisplatin-treated samples (GLMM fold change = 2.18, p-adj = 5.99 x 10⁻^6^) relative to untreated (Figure 2D, Extended Data Figure 1). To confirm this is not solely a feature of *TP53* deficiency, we conducted the same treatment and scWGS in 184-hTERT-L9 wild type cells, which showed similar drug-related patterns (Extended Data Figure 1, Supplementary Table 2). However, an investigation of event rates and mutational signatures on background copy number events from pseudobulk wild type and *TP53* deficient hTERTs did not consistently distinguish between CX5461-treated and untreated samples (Supplementary Figure 3, Methods). These data provide preliminary evidence that acute-drug treatment increases mutational burden, detectable by scWGS in the foreground cell-wise difference layer and not necessarily in the background bulk layer.

To enable systematic quantification of foreground mutational events (Figure 1C), we developed a feature space to classify CNAs by key structural characteristics (Figure 1D; Extended Data Figure 2, Supplementary Table 3, Supplementary Table 4, Methods). We explored five uncrossed feature classes using random-forest classifiers to distinguish treated from untreated cells. Classification performance varied across experimental groups (held-out ROC–AUC = 0.62–0.96), and Gini importance and SHAP values were used to assess the contribution of individual features (Extended Data Figure 2, Supplementary Table 3, Supplementary Table 4, Methods). Based on this analysis and emphasizing biological interpretability, we settled on three key components: (1) size (4, 8], (8, 17], (17, 38] and >38 Mb; (2) chromosomal topology (telomere-bounded, centromere-bounded, interstitial, or whole arm); and (3) the direction of CN change relative to the cell’s MRCA (gain or loss). We used these classes to define a feature space composed of 32 mutually exclusive categories (4 sizes x 4 chromosomal topologies x 2 CN directions); ie each foreground CNA is assigned to only one feature (Figure 1D). We note that < 8 Mb whole-arm events do not exist in the human genome and were thus omitted from the feature space, leaving us with a total of 30 informative features. We characterised each cell’s foreground in this 30-feature space by enumerating the observed foreground events per cell for each feature class (Figure 1E). We observed broadly that several of these features separated drug-treated and untreated *TP53*^-/-^ and WT hTERTs (Figure 2E, Extended Data Figure 1, Supplementary Table 5). For example, large (17-38 Mb), telomerically-bounded losses (Figure 2E) were highly enriched in CX5461 (GLMM fold change = 3.79, *p-adj* = 1.84 x 10^-12^) and cisplatin (GLMM fold change = 3.87, *p-adj* = 3.06 x 10^-13^) cells. Taken together, these results show that a single cell genome sequence feature space capturing cell-wise events is able to identify drug induced mutations.

We reasoned that different generative processes likely contribute to patterns of foreground CNAs. Thus, we explored the use of a hierarchical Dirichlet process^31–35^ (HDP) for deconvoluting the extent of their contributions to the feature classes (Figure 1F). We fit a single HDP signal decomposition model^31^ over 43,516 single cell genomes across 163 samples (Table 1, Supplementary Table 6, Supplementary Table 7, Methods) from several experimental contrasts: including four cell lines and two PDX lines with treatment with 8 drugs along with untreated controls. The optimal solution (Supplementary Figure 4A,B, Supplementary Table 8, Methods) identified 8 composite mutational signatures across the entire dataset (Figure 1G, Figure 2F, Extended Data Figure 3). Signature similarity varied (max cosine similarity = 0.92, min cosine similarity = 0.03, Figure 2G). PCA on the signature loadings suggested that no single feature drove separation, and that the greatest source of separation was the gain-loss dichotomy (PC1 explained variance = 42%). Following signature discovery, cellular exposure to each signature is evaluated along with a measure of certainty ranging from 0 to 0.95 (Supplementary Figure 4C, Methods). Certainties of signature exposures moderately correlated with exposure (Spearman’s R 0.55, *p-adj* < 2.2 x 10^-16^, Supplementary Figure 4D), and mutational burden (Spearman’s R 0.32 to 0.56, *p-adj* < 2.2 x 10^-16^). Certainties did not correlate with feature coverage (Spearman’s R -0.08 to 0.28), indicating that confident signature assignment coincides with CNAs piling up in features as opposed to being spread across the feature space (Supplementary Figure 4E). For comparison of signature extraction, we also performed signal decomposition with non-negative matrix factorization (NMF), which was less discriminative. NMF found only four signatures (Supplementary Figure 5), all of which appeared similar to one or more signatures identified by HDP (cosine similarity range 0.83 to 0.95), but could not fully resolve the differences in signature exposure among experimental contrasts. Following foreground CNA mutational signature discovery, we next quantified how cellular exposure to signatures change across experimental contrasts in our data (Figure 1H).

**Table 1.** Summary of the data used in this study.

| Data set | Drug | N. samples | N. high quality cells | N. analysed cells |
| --- | --- | --- | --- | --- |
| **184-hTERT-L9 TP53-/-** | UNTREATED | 32 | 12458 | 3458 |
|  | CX5461 | 17 | 3838 | 2119 |
|  | CISPLATIN | 9 | 4159 | 2213 |
|  | PDS | 4 | 843 | 369 |
|  | PALBOCICLIB | 4 | 1620 | 465 |
| **Total 184-hTERT-L9 TP53-/-** |  | 66 | 22918 | 8624 |
| **184-hTERT-L9 Wild Type** | UNTREATED | 4 | 1982 | 341 |
|  | CX5461 | 5 | 1432 | 558 |
|  | CISPLATIN | 3 | 1532 | 439 |
|  | PDS | 1 | 721 | 232 |
| **Total 184-hTERT-L9 Wild Type** |  | 13 | 5667 | 1570 |
| **184-hTERT-L9 TP53-/- BRCA1-/-** | UNTREATED | 6 | 3816 | 2959 |
|  | CX5461 | 3 | 2053 | 1776 |
|  | CISPLATIN | 3 | 1536 | 1303 |
|  | OLAPARIB | 1 | 231 | 156 |
|  | PALBOCICLIB | 1 | 970 | 694 |
| **Total 184-hTERT-L9 TP53-/- BRCA1-/-** |  | 14 | 8606 | 6888 |
| **HEK-293 Wild Type** | UNTREATED | 2 | 1436 | 1110 |
|  | CX5461 | 1 | 553 | 402 |
|  | VORELOXIN | 2 | 917 | 693 |
|  | ETOPOSIDE | 2 | 598 | 459 |
|  | ZM447439 | 1 | 516 | 418 |
| **Total HEK-293 Wild Type** |  | 8 | 4020 | 3082 |
| **TNBC PDX SA609** | UNTREATED | 13 | 7886 | 5413 |
|  | CISPLATIN | 22 | 13069 | 10129 |
| **Total TNBC PDX SA609** |  | 35 | 20955 | 15542 |
| **TNBC PDX SA535** | UNTREATED | 6 | 2223 | 1703 |
|  | CX5461 | 10 | 3854 | 3057 |
|  | CISPLATIN | 11 | 5040 | 3050 |
| **Total TNBC PDX SA535** |  | 27 | 11117 | 7810 |
| **Entire data set** |  | 163 | 73283 | 43516 |

Columns from left to right: the genotype and cell of origin, treatment, number of samples, number of cells in all the row samples together after quality control and S-phase removal (see Methods), and number of cells used for HDP training (cells with $\leq 2$ foreground events are removed). The cisplatin and untreated PDX samples were previously published (Salehi et al, 2021). All other samples are new to this study.

### Mechanistically distinct DNA strand break-inducing drugs generate a telomere-bounded CNA signature in *TP53* deficient cells

We investigated whether drugs that induce DNA breaks by different DNA interacting mechanisms result in distinct foreground CNA patterns again using concentrations that induce a biological response without killing all cells (Supplementary Figure 1A). We used GLMMs (Methods) to quantify how cell-level mutational signature exposure changed in response to drug treatment. In isogenic *TP53*^-/-^ hTERT cells, we observed that treatment with either CX5461 or cisplatin converged on the same HDP-identified composite mutational signature (Figure 3A-C, Extended Data Figure 4), labelled as TboundedArm, as it is defined primarily by large (17-38 and >38 Mb) telomerically-bounded losses, and whole-arm losses (Figure 2F). Treatment of cells with each of these two drugs results in a significantly enriched cell-wise exposure (which represents the number of CNA events assigned to a signature) to the TboundedArm signature (CX5461: fold change = 11.5, *p-adj* = 3.07 x 10^-4^; cisplatin: fold change = 10.02, *p-adj* = 0.035), compared to untreated cells (mean exposure = 0.34). Treatment of cells with PDS – another G-quadruplex stabilizer structurally unrelated to CX5461, also converges on the TboundedArm signature (mean exposure = 2.38, N samples = 1, N cells = 147, Figure 3A-C).

**Figure 3.**
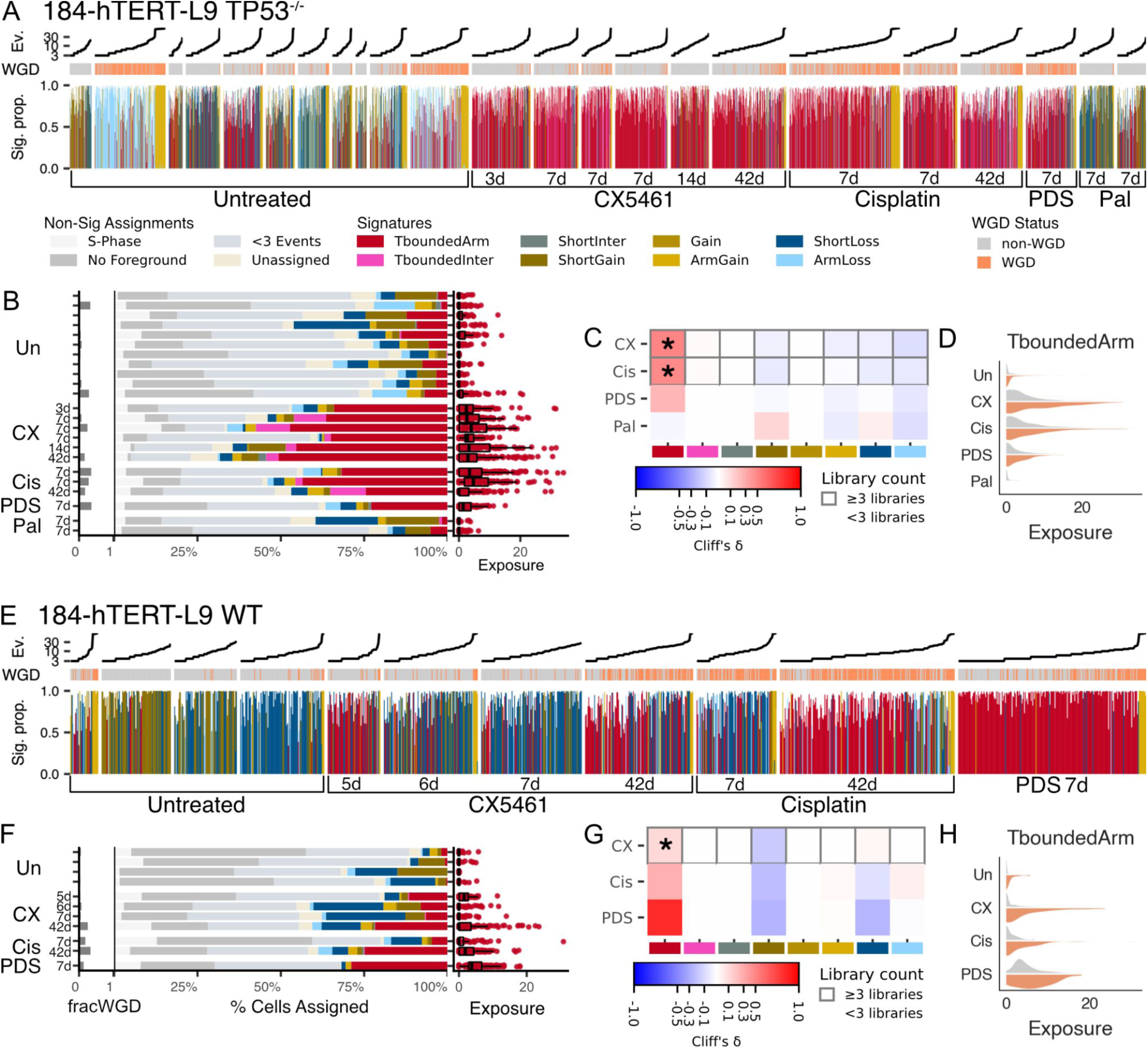
Foreground mutational signatures in HR-proficient 184-hTERT-L9 cell lines. *TP53*-deficient 184-hTERT-L9 cells were subjected to a panel of drug conditions, then processed by foreground extraction and HDP signature decomposition. **A,** Each column shows the per-cell percent exposure to each signature. Events with <0.5 certainty in assignment are included in the percentage calculation but plotted as white space. Cell-wise event rate and whole-genome doubling (WGD) status are annotated above the plot. **B,** Cells are assigned to their maximally exposed signature. Proportion of cells assigned to each signature, or the reason they were filtered out prior to signature decomposition. The left margin shows the fraction of WGD cells per library; the right margin shows cell-wise exposure to the drug signature (TboundedArm) per library. **C,** Summary of changes in exposure following drug treatment across all signatures. Grey boxes mark drugs with sufficient sample size for generalized linear mixed model (GLMM) comparison; stars denote statistical significance (*p-adj* < 0.05); color encodes effect size (Cliff’s Delta). **D,** Summary of differences in TboundedArm exposure between WGD and non-WGD cells. **E–H,** As in A–D, but for wild-type 184-hTERT-L9 samples.

We next asked whether DNA replication states or whole genome duplication intersects with drug induced mutation signatures. We first examined whether S-phase cells could account for the presence of TboundedArm. As we and others have shown, DLP+ sequencing can identify cells in S-phase^9,36,37^ due to variations in copy number in early and late replicating regions of the genome. HDP signature decomposition resolved two signatures – ShortGain and ShortLoss – that were enriched in short interstitial gains or losses respectively (Figure 2F, Extended Data Figure 3), and were found throughout samples of WT and *TP53*^-/-^ cells regardless of treatment (Figure 3A,E). We hypothesized that S-phase cells^9,37^ that were missed during quality control (Methods) could account for these signatures. To test this, we isolated 77 wild type untreated cells identified as S-phase^9^, held the 8 signatures fixed, and estimated each S-phase cell’s exposure to the signatures via maximum a posteriori (MAP) estimation under a multinomial mixture model. S-phase cells were commonly exposed to ShortGain (mean exposure = 5.98) or ShortLoss (mean exposure = 3.09), and were not exposed to TboundedArm (mean exposure = 0.05). This indicates that S-phase cells were unlikely to be responsible for TboundedArm discovery, and furthermore, insofar as they slip through quality control measures, they would segregate into ShortGain or ShortLoss. To corroborate this in our training data, we looked at exposures in drug-treated samples, where we reasoned that cellular exposure to TboundedArm should not coincide with exposure to the Short signatures. Indeed, exposures were not positively correlated (for example CX5461-treated *TP53*^-/-^: ShortGain and TboundedArm Spearman R = -0.28; ShortLoss and TboundedArm Spearman R = -0.28) – indicating that the process that leads to TboundedArm did not co-exist with the process that leads to ShortGain or ShortLoss. Together these results suggest that CNA patterns from single S-phase cells do not explain the drug-related telomere-bounded signature.

Cell cycle arrest may also be a consequence of DNA damage induction. We investigated next whether non-mutagen induced cell cycle arrest results in the telomere-bounded events. We treated *TP53*^-/-^ hTERT cells with IC30 palbociclib, a CDK4/6 inhibitor (Palb, 29 nM), at a concentration sufficient to induce G1 checkpoint arrest (Extended Data Figure 5A,B) without causing widespread DNA damage (Extended Data Figure 5C). The non-DNA damaging drug (Figure 3A-D) did not induce the drug-associated TboundedArm signature (mean exposure = 0.26, N samples = 2, N cells = 219).

Finally, replication leading to whole-genome duplication (WGD) is prominent across human cancers and is implicated in tumour progression: including the development of resistance, metastasis, and instability^30,38–41^. Given that cells with WGD contain more genomic substrate, we reasoned that drug-treated cells with WGD might exhibit more exposure (which again reflects the number of events assigned to a signature) to the TboundedArm signature than without-WGD cells. In CX5461- and cisplatin-treated *TP53*^-/-^ hTERTs, we found that cells that had undergone WGD had increased exposure to the TboundedArm signature (Figure 3D) compared to non-WGD cells (CX5461: fold change = 1.86, p-adj = 2.22× 10^-9^, cisplatin: fold change = 1.46, p-adj = 2.45 × 10^-4^). PDS-treated samples (N samples = 1, N cells = 147) also showed this trend (fold change = 1.48). We note that we did not find a signature specific to WGD status, thus the background WGD state appears to influence the prevalence but not the form of the mutational processes involved.

Together, these results affirm that TboundedArm is a drug signature associated with DNA-interacting small molecules that create double strand breaks by different mechanisms which is increased in WGD backgrounds but appears unrelated in origin to S-phase or cell cycle arrested states.

### Influence of DNA repair and active genomic instability on drug induced mutational signatures

The consequences of drug induced strand breaks may vary in relation to somatic mutations affecting DNA damage sensing or DNA repair, as well as cell lineage. Thus, we next examined the signature exposure in isogenic cell lines with different gene knockouts for damage sensing and repair. We reasoned that as *TP53* deficiency, which is highly prevalent in breast cancers^42^, enables progression through TP53-dependent DNA damage checkpoints^43,44^, we may observe exacerbated signature exposure compared to TP53-proficient cells. To calibrate this understanding, we looked to wild-type hTERTs, where we observed high exposure to the TboundedArm signature (Figure 3E-H) in CX5461-treated cells (fold change = 9.04, p-adj = 0.049) compared to untreated (mean exposure = 0.07, N samples = 4, N cells = 341); we saw similar trends in cisplatin-treated (fold change = 17.01, N samples = 2, N cells = 331) and PDS-treated samples (fold change = 37.25, N samples = 1, N cells = 232). Noting convergence on TboundedArm as a drug signature in both WT and *TP53*^-/-^ hTERTs, we next compared their exposures to drug-induced TboundedArm. Compared to WT CX5461-treated cells, we found that the *TP53*^-/-^ CX5461-treated cells (Figure 3A,E) had significantly higher exposure (fold change = 4.19, *p-adj =* 6.03 x 10^-3^), consistent with *TP53* deficiency enabling progress through TP53-dependent DNA damage-sensing checkpoints^43,44^. Furthermore, when we assign cells to their highest exposed signature (Figure 3B,F, Methods), we found that a greater proportion of *TP53*^-/-^ cells were assigned to TboundedArm (40%) compared to WT cells (12%), indicating that *TP53*^-/-^ cells were more likely to be affected by treatment. We saw the same patterns in cisplatin treated samples (fold change = 2.00, 15% → 32% cells assigned), but not in the smaller sample sets (232 WT vs 147 *TP53*^-/-^ cells) of PDS exposed cells (fold change = 0.42, 29% → 23% cells assigned). Taken together, we found that for at least two drugs, *TP53* deficiency is permissive to the detection of the TboundedArm signature. However there may be drug-dependent differences in DNA damage sensing with pyridostatin.

DNA repair processes influence the outcome of drug action on the genome, and triple negative breast cancers commonly inactivate the homologous recombination repair pathway (i.e., HR-deficiency, HRD)^45^, which renders cells sensitive to DSB-inducing drugs such as cisplatin^46^ and CX5461^13^. To address this, we performed foreground mutational signature analysis on treated and untreated isogenic *TP53*/*BRCA1*-inactivated mammary 184-hTERT-L9 cells *(BRCA1*^-/-^ hTERTs, see Supplementary Figure 1C,D for IC30). We noted that untreated cells already carried a high total cell-wise mutational event rate (66.38 CNAs/cell, Extended Data Figure 1), and foreground event rate (11.96 CNAs/cell), consistent with ongoing genomic instability. Although *BRCA1*^-/-^ cells are more sensitive to CX5461 and cisplatin (Supplementary Figure 1C,D), we noted no measurable difference in foreground event rate from CX5461 treatment (fold change = 1.35, p-adj = 0.18), and minimal biological effect from cisplatin treatment (fold change = 1.59, p-adj = 4.97 × 10⁻^3^). Signature decomposition classified the cell line’s endogenous genomic instability as a distinct, but telomere bounded signature class: TboundedInter (Figure 2F, Figure 4A-D, Extended Data Figure 6, mean exposure = 7.36). Endogenous TboundedInter is similar to TboundedArm (cosine similarity = 0.81), sharing large telomerically-bounded gains and losses — which notably are a minority component of other signature classes (Figure 2F, Extended Data Figure 3). It differs from the TboundedArm signature in that it contains small interstitial gains and losses, and lacks large arm losses. Surprisingly, when we treated *BRCA1*^-/-^ hTERTs with CX5461 or cisplatin we did not detect exposure to TboundedArm (Figure 4A-C, mean exposure = 0.52, 0.52 respectively), nor was a new signature detected. We also did not observe a meaningful increase in exposure to the TboundedInter signature (fold change = 1.4, *p-ad*j = 1; fold change = 1.2, *p-adj* = 1, respectively). As *BRCA1*-deficient cells are also sensitive to PARP inhibitors, we tested treatment with olaparib (Figure 4A-C). However, we also did not detect a new signature above the high endogenous telomere-bounded signature class (TboundedInter mean exposure = 11.2, N samples = 1, N cells = 156). Finally, consistent with previous results, treatment with palbociclib had no effect (TboundedInter mean exposure = 5.84, N samples = 1, N cells = 694, Figure 4A-C), and exposure to TboundedInter tended to be elevated in WGD cells (2.38 fold change over non-WGD cells, *p-adj =* 4.71 x 10^-170^, Figure 4D). Taken together, we conclude that 184-hTERT TP53/*BRCA1* mutant HR-deficient cells already possess a very high endogenous telomere bounded cell-wise mutational foreground, consistent with the role of HR in telomere maintenance^47^, which obscures observation of drug induced mutational events.

**Figure 4.**
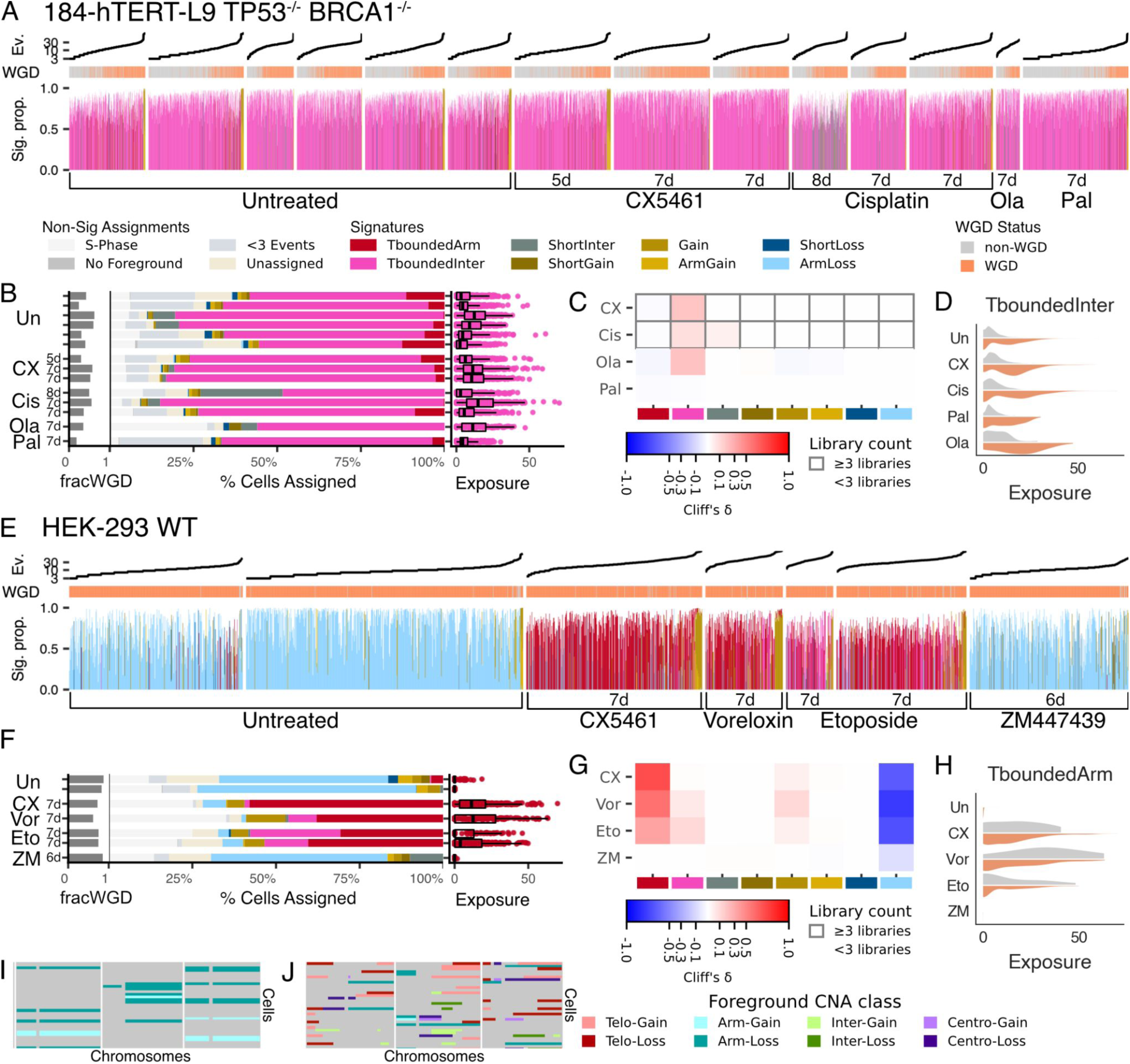
Foreground mutational signatures in large-scale genomically unstable cell lines. *TP53-BRCA1*-deficient 184-hTERT-L9 cells were subjected to a panel of drug conditions, then processed by foreground extraction and HDP signature decomposition. **A,** Each column shows the per-cell percent exposure to each signature. Events with <0.5 certainty in assignment are included in the percentage calculation but plotted as white space. Cell-wise event rate and whole-genome doubling (WGD) status are annotated above the plot. **B,** Cells are assigned to their maximally exposed signature. Proportion of cells assigned to each signature, or the reason they were filtered out prior to signature decomposition. The left margin shows the fraction of WGD cells per library; the right margin shows cell-wise exposure to the endogenous TboundedInter. **C,** Summary of changes in exposure following drug treatment across all signatures. Grey boxes mark drugs with sufficient sample size for generalized linear mixed model (GLMM) comparison; color encodes effect size (Cliff’s Delta). **D,** Summary of differences in TboundedInter exposure between WGD and non-WGD cells. **E–H,** As in A–D, but for HEK293 WT cells with TboundedArm. **I,** examples of foreground events in untreated and **(J)** CX5461-treated cells on chromosomes 4,5 and 6.

We next asked how non-HR related, high cell-to-cell chromosomal instability phenotypes interact with drug induced telomere bounded mutations. We conducted scWGS on HEK-293 cells transformed by adenovirus fragments containing the transcription units of E1A/E1B^48^, which exhibit genome instability and observed that this lineage exhibits foreground characterised by arm level or chromosome level losses. HEK-293 cells have dominant WGD (84%, Figure 4E,F,I, Extended Data Figure 7). We observed high exposure to a signature (Figure 2F, Extended Data Figure 3) that we call ArmLoss (Figure 4E–I, mean exposure = 5.86, N samples = 2, N cells = 1110), reflecting the genomic instability of the line. Treatment with CX5461 resulted in clearly increased exposure to the TboundedArm signature observed in mammary epithelial lineage cells (Figure 4E–H,J, mean exposure = 14.00, N samples = 1, N cells = 402).

Finally, we asked whether a third mechanistic class of DSB inducer — topoisomerase II inhibition — would also elicit TboundedArm in the HEK-293 lineage. We assayed cells with two topoisomerase II inhibitors that act by different mechanisms: the DNA-intercalating voreloxin, which produces more targeted DNA strand breaks, and etoposide, which does not intercalate DNA but leaves TOP2 covalently linked to DNA globally. Treatment with either drug produced the same foreground signature, TboundedArm (voreloxin: mean exposure = 16.16, N samples = 1, N cells = 182; etoposide: mean exposure = 9.16, N samples = 2, N cells = 459). Although the number of exposed mutated cells is relatively small, etoposide and voreloxin also demonstrated contribution of TboundedInter (mean exposure = 5.60, 5.23 respectively), in contrast with CX5461 (mean exposure = 0.63). To again control for cell cycle arrest, we examined the effect of a non-clastogenic cell cycle inhibitor. As HEK-293 are less sensitive to CDK4/6i due to the E1A/E1B protein action, we treated HEK-293 with the aurora kinase inhibitor ZM-447439, which prevents mitotic entry but does not directly induce DNA damage. HEK-293 cells treated with ZM-447439 did not display the TboundedArm signature (mean exposure = 0.00, N samples = 1, N cells = 418) and instead retained the endogenous ArmLoss signature at levels similar to untreated cells (mean exposure = 5.41). Taken together, we conclude that DSB inducing small molecules of different mechanistic classes induce similar telomere-bounded chromosomal scale mutations, also apparent in different cell lineage backgrounds.

### Foreground mutational signatures are genomic pharmacodynamic measurements

Having established TboundedArm as a detectable drug-induced signature in HR-proficient cells, we next examined whether the kinetics of drug activity over time could be measured. As previously highlighted, the magnitude of cellular exposure to a signature is a direct measurement of the abundance of that signature’s foreground CNA patterns in a cell. Thus, we hypothesized that increasing drug dose would coincide with greater DNA damage resulting in an increase in exposure to TboundedArm. To test this, we treated *TP53*^-/-^ hTERT cells with two doses of CX5461 (27 nM and 703 nM). As expected, we observed a dose dependent response in the magnitude of TboundedArm (Figure 5A,B, mean exposure: 0.75 → 3.41). We also observed that a greater proportion of cells were affected (14% → 35%). We observed similar drug dose-dependent effects with two other comparisons we measured: cisplatin (93 nM, 1144 nM, Figure 5A,B, mean exposure 0.82 → 6.38, 11% → 44%) in *TP53*^-/-^ hTERTs, and voreloxin (50 nM, 150 nM) in HEK-293 cells (Figure 5A,B, mean exposure 4.14 → 16.16, 29% → 38%). These results exemplify that foreground signatures capture dose-related changes in cellular mutational burden, both in severity per cell and in the fraction of cells affected.

**Figure 5.**
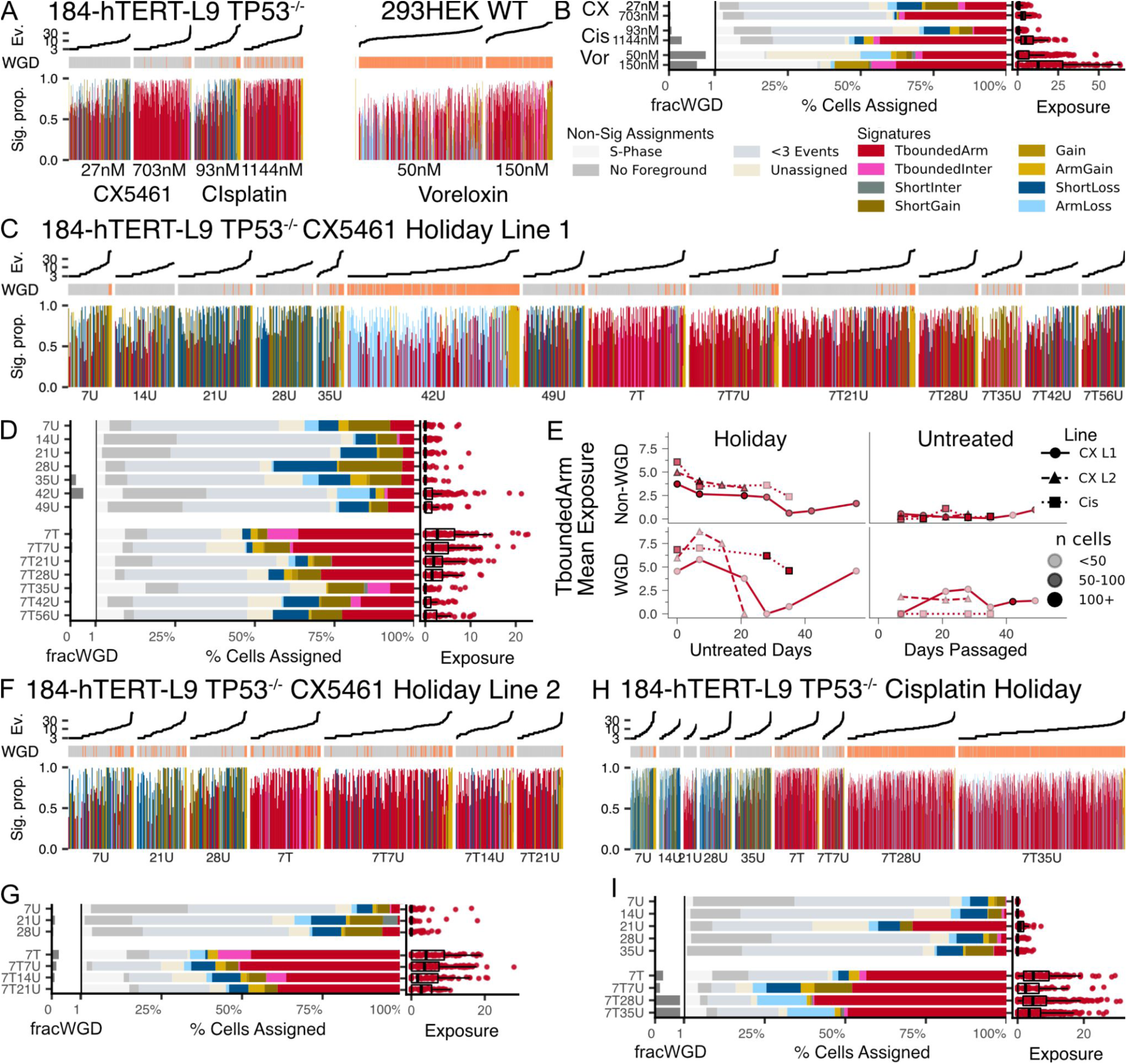
Foreground mutational signatures track drug pharmacodynamics. **A,** *TP53-*deficient 184-hTERT-L9 cells were treated with CX5461 or cisplatin at two doses; HEK293 cells with Voreloxin. Each column shows the per-cell percent exposure to each signature. Events with <0.5 certainty in assignment are included in the percentage calculation but plotted as white space. Cell-wise event rate and whole-genome doubling (WGD) status are annotated above the plot. **B,** Cells are assigned to their maximally exposed signature. Proportion of cells assigned to each signature, or the reason they were filtered out prior to signature decomposition. The left margin shows the fraction of WGD cells per library; the right margin shows cell-wise exposure to the drug signature (TboundedArm). **C,D,** *TP53*-deficient 184-hTERT-L9 were subjected to CX5461 for seven days, washed of drug, and then left untreated for up to 56 days (line 1). Matched untreated controls were also passaged. Sample annotations represent how many days of treatment (T), or days of lack of treatment (U) they were subjected to. **E,** Mean exposure to TboundedArm across all experiments, stratified by control vs holiday, and cell whole genome doubling status. Opacity encodes cells per sample. **F,G,** Replicate of C,D for 21 days of holiday (line 2). **H,I,** cisplatin treatment followed by 35 days of holiday.

Key to the action of DSB inducing small molecule drugs is the duration of action on the genome after treatment. We reasoned that the characteristic of foreground single cell mutational signatures to readout ongoing mutations should capture the mutational decay temporal dynamics. Hence, we measured the mutational foreground after drug wash-out, which we describe as a drug holiday following treatment. First, *TP53*^-/-^ hTERTs were treated with CX5461 for 7 days, after which the drug was withdrawn and the media changed every 7 days and cells passaged every 14 days in drug-free media (Methods). We measured single cell foreground mutational signatures every 7 days for up to 56 days (Figure 5C-G). As expected, exposure to the TboundedArm foreground signature declined progressively after the drug was washed out and throughout the holiday. Surprisingly, low-level residual signal remained detectable even up to the final holiday week (mean exposure = 1.7) – indicating that the mutational footprint of drug treatment persists in the genome for at least 3 weeks beyond the active treatment window. To determine whether persistence was specific to CX5461, we conducted similar holiday interval sampling after cisplatin exposure (Figure 5E,H,I, Supplementary Figure 6). We observed that late stage cisplatin holiday cells tended to have undergone WGD (28 holiday days = 86.7% WGD, 35 holiday days = 87.4% WGD). After controlling for exposure associated with WGD, cisplatin also showed a similar trend of signal decay (Figure 5E). Notably, among WGD cells, signal strongly persisted up to the last holiday sample (day 35). Taken together the data show that mutational persistence after treatment with different DSB inducing agents can last weeks after exposure.

### Foreground mutational signatures mirror drug resistance *in vivo*

Human tumours are clonally and mutationally heterogeneous^49^ and it is well-established that successive rounds of chemotherapy alter clonal composition^11,50,51^ and transcriptional states^51–53^, eventually selecting for resistant cells. Having shown that foreground signatures can track genomic pharmacodynamics in cell lines, we next asked whether they could be used to interrogate the emergence of drug resistance in vivo. A key question is whether resistance is accompanied by tolerance to ongoing mutations, or a reduced effect of agents in causing mutational burden. We monitored the foreground signature landscape of two PDX lines, both derived from invasive ductal breast carcinoma, across four transplant generations of a regimen designed to replicate chemotherapeutic drug holidays with cisplatin. As previously described^10,11,52,54^, each generation was transplanted into recipient mice, grown until 300 to 400 mm^3^, and then dosed; tumours were harvested once the tumour volume had shrunk by 50%, or reached 1000 mm^3^, homogenized, and split between DLP+ scWGS sequencing or transplantation into the next generation. Every treated generation was therefore dosed and sequenced, with the brief pre-treatment regrowth window constituting a *de facto* drug holiday. Alongside the continued treatment branch we propagated a matched untreated branch for each line^52^, and an extended drug holiday branch — where tumours from each treated generation were transplanted into mice and allowed to grow without drug dosing. Growth curves were recorded for every tumour, and growth status at the time of sequencing was classified according to modified RECIST guidelines^55^.

The first line we tested, SA609, harbours an intronic *BRCA1* mutation and a variant of unknown significance^11,52^ (see Supplementary Table 9 for tumour type status). Both first-generation treated samples (1T) were partially responsive (PR) to cisplatin and showed elevated absolute CNA burden (149.14 CNAs/cell, N samples = 2, N cells = 587) and foreground CNA burden (43.63 CNAs/cell) relative to their untreated counterpart (1U absolute = 124.68 CNAs/cell, foreground = 15.04 CNAs/cell, N samples = 2, N cells = 413; Figure 6A,B, Extended Data Figure 8). Signature decomposition revealed that SA609 1T cells were highly exposed to TboundedInter (Figure 6F,G,J; mean exposure = 34.31), confirming that cisplatin treatment increased a telomere-bounded foreground mutational signature, as in the *BRCA1*^-/-^ cell lines. In subsequent generations, as cisplatin responses shifted towards gradual resistance, with stable and then progressive disease, TboundedInter exposure fell (Figure 6E-G, mean exposure = 5.32 and 2.90, respectively) and was displaced by ShortInter (mean exposure = 1.87 → 5.50 → 7.48 respectively). ShortInter represents the background signature that dominates the untreated PDXs (all of which were classified as progressive disease; Figure 6C,D,K, mean exposure = 5.91). The two signatures are closely related (cosine similarity = 0.92), differing primarily in ShortInter’s heavier loading of short interstitial losses (Figure 2F, Extended Data Figure 3). In the extended holiday arms, tumours transplanted off the treatment branch and left undosed had reduced TboundedInter exposure relative to their treated parent generation (Figure 6H,I,L), lending further credence to the signature’s robustness.

**Figure 6.**
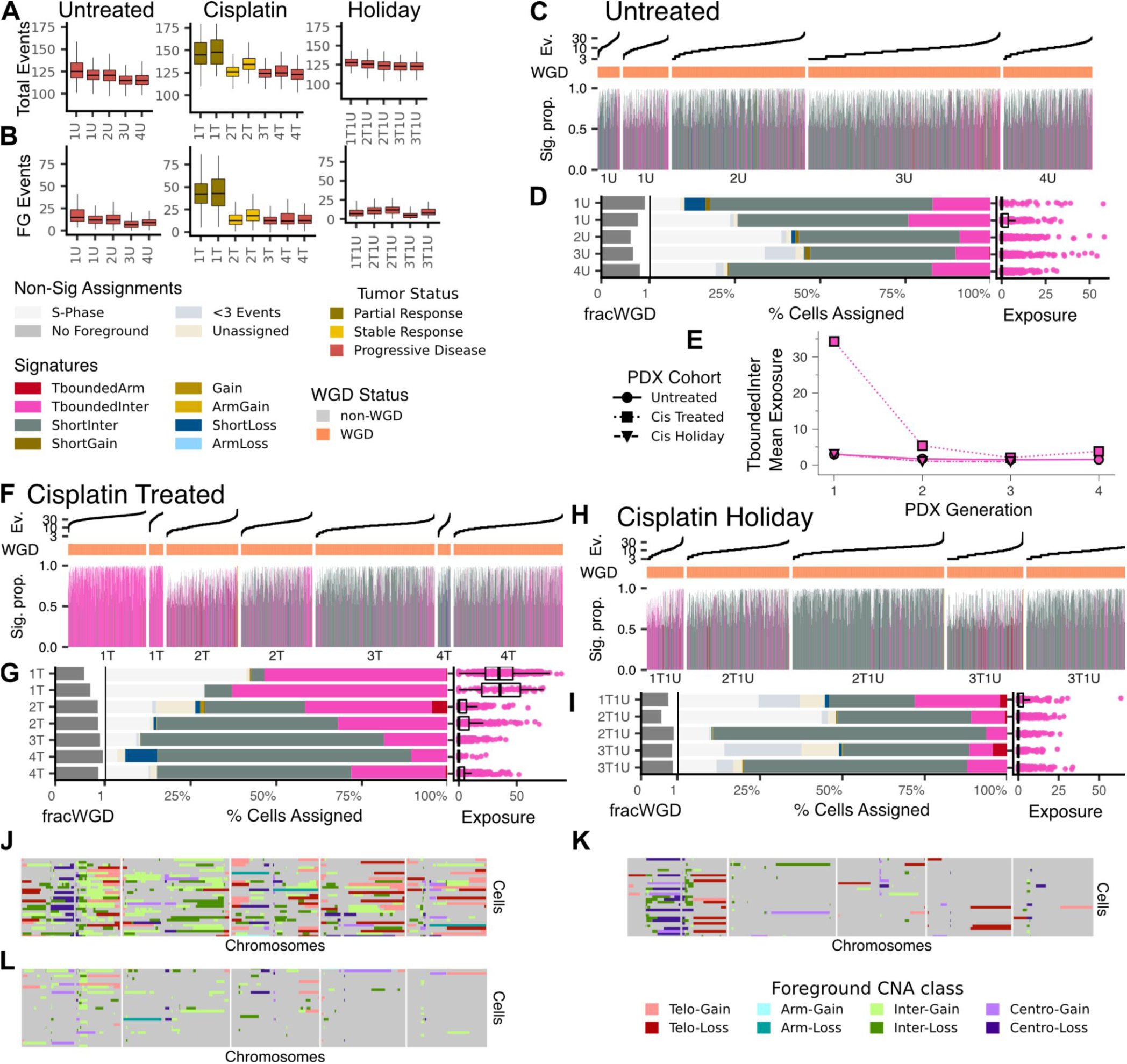
Foreground mutational signatures in SA609 PDX branches. SA609, a breast carcinoma PDX line, was propagated along three branches: treated with cisplatin at each transplant generation, left untreated across all generations, or treated and then given a drug-holiday generation. Generations are labeled with a number followed by T (treated) or U (untreated); for example, 1T is the first, treated generation. **A,** Event rates of absolute mutational CNAs for the three branches. Tumor status indicates response to drug at the time of sequencing. **B,** Foreground event rates. **C,** Per-cell proportional exposure to each signature in the untreated branch, with each column representing one cell. Events with <0.5 certainty in assignment are included in the percentage calculation but plotted as white space. Cell-wise event rate and whole-genome doubling (WGD) status are annotated above the plot. **D,** Cells are assigned to their maximally exposed signature. Proportion of cells assigned to each signature, or the reason they were filtered out prior to signature decomposition. The left margin shows the fraction of WGD cells per library; the right margin shows cell-wise exposure to TboundedInter. **E,** Summary of TboundedInter exposure across transplant generations for each branch. **F,G,** Exposures for the cisplatin-treated branch. **H,I,** Exposures for the cisplatin-holiday branch. **J,** Example foreground CN events on chromosomes 1 to 5 for cells highly exposed to TboundedInter 1T, (**K)** for cells highly exposed to ShortInter in the fourth treated generation (4T), and (**L)** for cells exposed to TboundedInter in the first holiday generation (1T1U).

Both of these patterns were broadly recapitulated in SA535^10^, which harbours a somatic *BRCA1* mutation^11,52^, under cisplatin and CX5461 treatment (Extended Data Figure 9). Despite differences in genotypic backgrounds, TboundedInter remained the drug-related signature in SA535, which once again peaked in the first treated generation of treatment (mean exposure: cisplatin = 5.12, CX5461 = 2.91), and declined as tumours progressed. However, both the initial signal and its subsequent decay were markedly attenuated relative to SA609, consistent with the weaker tumour growth inhibition observed in this model and with its previously reported clonal complexity and limited platinum sensitivity^11,52^. Together, these results show that cisplatin and CX5461 generate the same type of chromosomal scale mutational process *in vivo* and that onset of tumour resistance is associated with loss of the foreground mutational signature, implying loss of drug action on the genome, rather than tolerance to ongoing mutational events.

## Discussion

Strikingly, we observe that five different small molecules acting on DNA by three different mechanisms, cross linking of guanine bases, G-quadruplex associated DSB formation, and topoisomerase-2 inhibition, all result in a similar class of foreground large scale mutational event, identified as large gains and losses mapped to within 2.5 Mb of telomere regions. As short read sequencing cannot resolve telomeric low complexity repeat regions, we refer to these as telomere-bounded, however they may or may not include telomeric repeats. In support of this signature, drug-induced telomerically-involved alterations derived from DSBs are established effects of platinum cross-linkers^56,57^, G4-stabilizers^58,59^ and TOPII inhibitors^23,60^.

The mutational signatures induced by different DSB inducing agents used in cancer treatment are not well understood at genome scale as the mutations induced are not easily resolved by bulk WGS. Moreover, methods to clone and amplify single cells for mutational pattern analysis may be limited by the selective pressure of clonal amplification (Supplementary Figure 3). It is possible that the effects of some carcinogens may be entirely large, chromosomal scale mutations that preclude further cell division. The analytical procedure for mapping large scale mutational processes through DLP+ single cell genome sequencing used here has the advantage of being able to detect mutations above the scale limit of resolution of transposition based scWGS. Since all cells are sampled without a requirement for cell division, biases which would result from negatively selected mutations during clonal cell propagation after a mutational event are reduced. Under the hypothesis that cell-intrinsic foreground mutational processes and DNA replication should have a different form to drug induced mutational processes, hierarchical Dirichlet process decomposition identifies clearly separable signatures of DNA replication, endogenous processes and drug-associated signatures.

The mutational signatures induced are detectable in different cell type backgrounds with different states of DNA damage sensing (*TP53* WT and *TP53* deficient) and in the context of existing chromosomal instability processes (HEK-293). We observed that an isogenic *BRCA1* knockout breast epithelial cell line exhibited a high endogenous (i.e., without drug treatment) cell-to-cell mutational foreground with a telomere-bounded signature closely resembling that of drug treated *BRCA1* wildtype cells. The high cell intrinsic mutational exposure of *BRCA1^-/-^* cells may be consistent with the role of homologous recombination in telomere maintenance, reflected in telomere loss that we and others have reported cytogenetically. We could not measure a strong drug-related effect in these cells. The already high level of endogenous cell to cell foreground genomic instability of the same signature type as non HRD cells likely obscures the effects of DSB inducing drugs. Also the lower absolute drug exposure at IC30 level, balancing the need to retrieve sufficient cells for analysis, may reduce observable new mutations. It may also simply reflect very high levels of ongoing unrepaired, or erroneously repaired double strand breaks across the genome.

The limitations of phylogenetic reconstruction for large scale copy number single cell genomes are still an open problem despite intensive work on different computational approaches. In an MRCA based approach, foreground extraction of each cell is dependent on accurate phylogenetic reconstruction and MRCA inference; it is undeniable that different algorithms will result in various phylogenetic reconstructions and hence may affect mutational signature discovery, and cellular assignments to exposures^61–63^. In particular, genomically unstable lines have higher cell-to-cell variance, enfeebling phylogenetic reconstruction, increasing the average distance between leaf-cells and their MRCA, and thereby possibly undermining the quality and interpretability of foreground extraction. Despite this, our signature calls remained stable across the breadth of decomposition uncertainty thresholds we examined (Supplementary Figure 7), suggesting they are broadly consistent. Mutational signatures will in any case be dominated by the numerically greater mutational variations captured in single cells at the leaves of a tree rather than internal nodes.

The drug dose-dependent nature of the foreground mutational signatures admits decoding drug pharmacodynamics in terms of genomic effects. The induction of DSBs with the 5 agents studied is normally detectable within a few hours of exposure by DNA damage focus assays. However, DNA repair switches off after a few days or if cells arrest, precluding study of the temporal decay of mutational events following drug withdrawal. Remarkably, the drug induced foreground signatures from cisplatin and CX5461 were detectable in diploid cell populations for at least 3 weeks after drug removal. Both drugs induce DSBs but whereas cisplatin induces covalent cross-links, CX5461 is thought to interact in part via G-quadruplex secondary structures non-covalently. A further observation is that the cisplatin treated state was associated with dominant WGD in independent experiments. In WGD cells, the decay of telomere-bounded events appeared to be much slower, which may be related to slower replication and/or greater buffering capacity for mutations. The mechanism of persistence of the foreground is unknown. It could represent low levels of drug persistence in the cell, or cells exhibiting ongoing damage at replication, or cells that have dropped out of cycle with mutations but persist until they are diluted out by replication of non-mutated cells.

In TNBC patient derived xenografts treated with cisplatin or CX5461, we observed the same telomere-bounded mutational signatures as in *BRCA1*-deficient cell cultures, suggesting these mutational patterns are observed in tissues. We asked whether induced drug resistance in this setting is associated with tolerance to mutations or a reducing genomic mutational impact of the drug. In the former case mutations would still be observed above the background, whereas in the latter case mutational exposure should decline as tumours become more resistant. Under repeated cycles of cisplatin or CX5461, partially responsive tumours accumulate telomeric-bounded copy-number patterns (TboundedInter). These PDX-bearing mice were treated with 1/3 maximum tolerated dose (similar rationale to the use of IC30 in cell lines) of the drugs which would cause some killing in drug-susceptible clones of the PDX while retaining sufficient sample for analysis thereby eliciting a partial tumour response^11^. We were able to measure a foreground mutational signature from these initially responsive PDX lines. However, subsequent passages of these PDX lines displayed decreasing sensitivity and ultimately resistance to cisplatin or CX5461. Passages exhibiting low or no tumour responses exhibited decreasing exposure of the drug-associated mutational signature. After initial treatment of cisplatin and a first drug holiday (passage without cisplatin) for a single generation, the initial drug-associated mutational signal partially reverts towards pre-treatment levels. Subsequent passages that were fully drug resistant exhibited close to background levels only. Taken together this suggests that the mechanism of induced resistance in these cases results from reduced drug action on the genome, rather than tolerance to ongoing mutations.

Our results show that scaled single cell genome sequencing can detect cell-private mutational processes that are not easily detectable by other means. We suggest this approach may be used in the future to study environmental carcinogens and other potentially mutagenic molecules that may induce chromosomal-scale mutation rather than single base mutations.

## Methods

### Experimental Methods

#### Human cell lines

HEK-293 (adenovirus-transformed human embryonic kidney cell line^64^) is grown in DMEM/F-12 (Gibco), supplemented by 10% fetal bovine serum (FBS). The wild-type *hTERT*-immortalized non-transformed human mammary epithelial cell line 184-hTERT-L9^65^ and isogenic derivatives (*TP53*^−/−^, *TP53*^−/−^*BRCA1*^−/−^)^10^ were cultured as previously described in mammary epithelial basal medium (MEBM; Lonza) supplemented with the SingleQuots additives (Lonza), 5 μg/ml transferrin (Sigma-Aldrich) and 10 μM isoproterenol (Sigma-Aldrich). Cell lines were routinely confirmed mycoplasma-free and authenticated by STR profiling and allele genotyping (hTERT isogenics).

#### Chemicals

The following compounds were used in this study: CX5461, obtained either from Selleckchem (cat. no. S2684) or from Senhwa Biosciences as pidnarulex; cisplatin, supplied as cisplatin Injection BP (1 mg/mL; DIN 02355183; Accord Healthcare Inc.); etoposide (Sigma-Aldrich, cat. no. E1383); olaparib (AZD2281, Ku-0059436) (Selleck Chemical, cat. no. S1060); palbociclib (Selleckchem, cat. no. S1116); pyridostatin hydrochloride (PDS·HCl), provided as a gift by S. Balasubramanian (University of Cambridge, Cambridge, UK); voreloxin (Selleckchem, cat. no. S7518); and ZM-447439 (Cedarlane Labs, cat. no. A11009-5).

#### IC30 Determination

Cell lines with genomic knockouts were treated at partially inhibitory but recoverable drug concentrations, generally around IC30. IC30 was measured as the concentration at which 70% of cells remained viable in crystal violet growth assays, or, by a 30% decrease in metabolic activity of cells compared to the initial untreated condition (Cellular metabolic activity was assessed by measuring intracellular ATP levels using the CellTiter-Glo® Luminescent Cell Viability Assay (Promega)). The exact treatment doses (Supplementary Table 1) and experimental endpoints are labelled in the corresponding figures.

#### Drug Treatment and Harvesting for DLP+

Generally, cells were seeded in six-well plates and the following day, 1-2 ml of fresh media containing the given concentration of drug was added for the respective treatment time (e.g. 3-7 days) before the cells were washed two to three times with PBS to remove the drug and the single cells from replica-treated wells were pooled and harvested for DLP+ by trypsinisation. Cells from parallel wells were counted using a haemocytometer or subjected to flow cytometric analysis after fixation with 70% ethanol and staining with DAPI. For drug withdrawal (“Drug holiday”) experiments, two parallel plates were treated in duplicate for 7 days with the drug, before being washed off. One plate was harvested by pooling several wells of the same condition for single-cell DNA sequencing. The remaining parallel plate received fresh drug-free media and was incubated for a further 7 days for a total of 14 days before splitting. Cells were maintained under drug-free conditions by changing the media every 7 days and serially passaging every 14 days. The untreated cells were passaged in parallel as time-matched controls. At each time point, cells from three to six replicate wells were pooled and collected for DLP+ single-cell whole-genome sequencing, and the remaining cells were reseeded to generate the subsequent recovery time point.

#### Immunofluorescence in cell lines

Cells were stained for DNA damage markers in a similar manner as previously described by Xu et al.^13^. Briefly, hTERT *TP53*-/- cells were seeded at 20,000 cells per well of an 8-well chamber slide (Nunc™ Lab-Tek™ II Chamber Slide™ System Cat #154453, Thermo Scientific or Millicell® EZ Slide, Cat #PEZGS0816, EMD Millipore). Each well was a separate condition. Cells were treated at the respective concentration of the drug for the given time (2-48 hr). At the stated time, the media was removed and the cells were washed with PBS before fixing with 1.2% formaldehyde for 10 min at room temperature and permeabilisation with 0.01% Triton X-100 in PBS for 10 min, then washed and blocked with 4% BSA/PBS for 10 min. The cells were incubated with or without the primary and secondary antibodies. γ-H2AX was detected by 1/800 rabbit anti-H2A.X (phospho S139) antibody [EP854(2)Y] (Abcam #ab81299) and 1/2000 Goat anti-Rabbit IgG (H+L) Cross-Adsorbed Secondary Antibody, Rhodamine Red™-X (Invitrogen #R-6394) and 53BP1 was detected by 1/500 mouse anti-53BP1 Antibody, clone BP13 (EMD-Millipore #MAB3802) and 1/2400 Invitrogen™ Goat anti-Mouse IgG (H+L) Cross-Adsorbed Secondary Antibody, Alexa Fluor™ 488 (Fisher Scientific #A11001) and 20 µg/ml DAPI was used to stain the DNA for nuclear segmentation. The slide was then mounted with anti-fade (Vectashield) then stored in the dark at 4°C until ready for imaging. Images were acquired on a Zeiss LSM800 Confocal Microscope using a 20x objective and the images were subjected to nuclear segmentation, nuclear foci identification and co-localisation using an automated custom python script.

#### Serial passaging of PDX

All SA609 and SA535 untreated and cisplatin genomes were previously published^10,11,52^, with SA535 CX5461 treatment presented here for the first time. In brief, for serial passaging of PDX, xenograft-bearing mice were euthanized when the size of the tumours approached 1,000 mm^3^ in volume (combining together the sizes of individual tumours when more than one was present). The tumour material was excised aseptically, and processed as described for primary tumours. In brief, the tumour was harvested and minced finely with scalpels then mechanically disaggregated for one minute using a Stomacher 80 Biomaster (Seward) in 1 ml to 2 ml cold DMEM-F12 medium with glucose, l-glutamine and HEPES. Aliquots from the resulting suspension of cells and fragments were used for xenotransplants in the next generation of mice and cryopreserved. Serially transplanted aliquots represented approximately 0.1–0.3% of the original tumour volume. Treated and untreated TNBC-SA609 and TNBC-SA535 PDX were passaged up to 10 generations and scWGS was carried out at each time point. We annotate PDX samples based on their history, with “T”s indicating a treated generation, “U”s indicating an untreated generation, and prepended integers indicating the number of generations of that type. For example, 2T1U indicates a sample from a branch that was treated in two generations, with the final generation being untreated. In this paper, for clarity, we dropped generations that occurred prior to treatment from our naming convention so as to avoid consistent prepended “U”s. The SA535 time series was generated in the same way for 4–5 passages. Due to failed sequencing of the SA535 1T that was propagated, we used a 1T from a different treated branch.

#### PDX germline and somatic annotation

The genomic backgrounds and acquired mutations of PDX samples were assessed to identify sample HRD status, identify possible therapeutic vulnerabilities, and investigate the presence of any reversion mutations. We assessed the germline SNPs and somatic short mutations (SNVs and indels) in each PDX line. Briefly, bulk WGS sequencing data from patient tumour samples used to establish the PDX lines and patient matched normal were mapped to the GRCh38 reference genome following GATK Best Practices workflow^66,67^. Bulk and pseudobulk WGS data from PDX samples were first filtered to exclude mouse reads by aligning to a mm10 and hg38 concatenated reference genome. The remaining reads were then processed using the same workflow as tumour samples. Germline variants were called in the matched normal samples using HaplotypeCaller^68^ and annotated with SnpEff^69^ to provide genomic context and predict functional impacts, and cross-referenced against dbSNP, gnomAD and ClinVar^70^ to obtain population allele frequencies and clinical significance. Somatic variants were called using Mutect2^71^ in the tumour-normal setting. The identified short variants were then filtered to remove artifacts, technical noise, and likely germline mutations with FilterMutectCall and FilterAlignmentArtifacts (version 4.1.3.0). Filtered variants were next genomically and functionally annotated using SnpEff and GATK Funcotator, cross-referencing against the COSMIC^72^, ClinVar, and Gencode databases.

#### Ethics Approvals

All animal experiments were approved by the University of British Columbia Animal Care Committee under protocols A19-0298 and A24-0020. Patient tissue collection and establishment of SA609 and SA535 PDX lines were conducted with written informed consent under protocols approved by the University of British Columbia / BC Cancer Research Ethics Board (REB protocols H16-01625 and H20-00170).

### Single Cell DNA Sequencing and Analysis

#### Library preparation and sequencing

Cells in solution were prepared for single-cell DNA whole-genome sequencing following a previously described protocol^9^. Briefly, cells were dispensed into Wafergen nanowell chips pre-loaded with Illumina-compatible dual-barcoded indices with the Cellenion CellenONE piezoelectric dispenser. A one-pot reaction dissociates the cells within the well, exposing the genomic DNA allowing the Tn5 transposons in solution to randomly insert and fragment the genomic DNA with PCR primers and one of two barcoded indices. An on-chip PCR cycle then amplifies fragments with proper PCR primer combinations. The contents of all wells are eluted and pooled by centrifugation, and subjected to clean-up and size selection, followed by standard Illumina WGS sequencing. The majority of sequencing was outsourced to Canada’s Michael Smith Genome Sciences Centre, from which we received chastity-filtered, high-quality BCL-converted FASTQ reads for each DLP+ library.

#### Pre-processing, alignment, and state calling

Prior to alignment, FASTQ reads are run through FastQ-Screen^73^ and flagged for matches to *Homo sapiens* (GRCh37-lite), *Mus musculus* (GRCm38), *Oncorhynchus kisutch* (GCF_002021735.1), and a custom collection of 371 species from the *Pseudomonas* taxa. After FASTQ reads are annotated, they are aligned as previously described^30^: using TrimGalore for pre-processing (https://github.com/FelixKrueger/TrimGalore), bwa-mem v0.7.17^74^ for short-read alignment and Picard for alignment metrics (http://broadinstitute.github.io/picard/).

To generate human state calls, aligned cells are passed to the Mondrian pipeline^9^ (https://github.com/mondrian-scwgs/mondrian) developed by our collaborators^30^. In brief, aligned reads are binned into 500 kb windows, the counts are normalized for each bins GC content, and CNAs are called using HMMcopy. The specific version of Mondrian we used is a fork (https://github.com/molonc/mondrian_nf and https://github.com/bpotvin-bccrc/mondrian_utils); the core functionality and workflow were identical, with the exception of the Pseudomonas contamination filter.

#### Quality control and cell filtering

We assessed cell state calling quality as previously described in Laks et al^9^. We leveraged the outputs of Picard alignments and HMMcopy to compute several metrics which were fed into two random forest algorithms, the first of which produced cell quality scores ranging from 0 to 1, and the second of which assigns a probability that the cell is in S-phase. To ensure high quality foreground extraction, we retained cells with a quality score of ≥ 0.75, and a probability of being in S-phase of < 0.5. We applied a new contamination metric using FastQ-Screen flags to estimate the number of cells that are eukaryotic in origin (i.e. human, mouse or salmon) and everything else assumed to be bacterial (i.e. pseudomonas, unknown). We dropped cells that have >65% non-eukaryotic reads. We then drop cells that have less than 60% human reads of total eukaryotic reads. This leaves us with high quality, predominantly non-S-phase, non-contaminated, human cells.

#### hTERT cell genomic identity confirmation

To verify the genotype of hTERT cell line samples and rule out cross-contamination, we analysed the single nucleotide polymorphism (SNP) mutations in each sample. First, quality-filtered single-cell read alignments, mapped to the GRCh37 reference genome, for each scWGS library were merged into a per-sample pseudobulk alignment to enable variant calling. We then used HaplotypeCaller^68^ to call SNPs, which were subsequently validated and annotated for SNPs using GATK’s ValidateVariants and snpEff^69^. The resulting consensus genotypes were compared against reference baseline profiles to confirm cell line identity and purity.

#### Bulk copy number alteration signatures in hTERT cells

To identify copy number alterations (CNA) events in the hTERT cells at the bulk level, we used Sequenza^75^ to resolve allele-specific copy number profiles and detect LOH events in the pseudobulk hTERT cell line data re-mapped to hg38. We implemented the WGS Sequenza workflow in matched tumour-normal mode, where an untreated L9.WT bulk WGS sample was designated as the matched normal sample for all pseudobulk samples, including the WT, *TP53*^-/-^ and *TP53*^-/-^BRCA1^-/-^ genotypes. The resulting segment files were used to extract CN signatures and compare sample CNA profiles using the sigminer R package^76^. To ensure robust CNA signature extraction, we implemented two complementary classification frameworks. We applied the 48-component framework by Steele et al.^29^, as well as the expanded 80-component framework by Wang et al.^76^ to account for structural transitions and maintain cross-condition stability. We used non-negative matrix factorization (NMF) for de novo extraction of signatures from hTERT CNA profiles decomposed into the 48-component or 80-component features. A maximum of 12 signatures were evaluated, with 50 NMF replicates, with the solution maximizing stability while minimizing mean cosine distance selected for final signatures, and NMF was rerun an additional 50 times with the selected number of solutions. Signature activities of the de novo CN signatures across our hTERT samples were calculated using quadratic programming (QP). Similarity of sample 48-component and 80-component profiles was evaluated by cosine similarity. Analyses and data visualization were performed using R 4.3.2.

### Foreground and MRCA Extraction

We established a computational framework that extracts allele-specific non-clonal CNA events from the clonal background of a population of cells. First, cells passing our quality thresholds are passed to SIGNALS^10^ (selftransitionprob = 0.99, firstpassfiltering = FALSE) to compute the allele-status of each CNA. To help reduce noise, non-confident short breakpoints are removed using a multi-sample piecewise constant fitting step, where cell allele-aware CN profiles are normalized within each library. Then, we provide each cell’s copy number state and its allelic status (allele A or B) to MEDICC2^62^ for the construction of allele-aware phylogenies. We leverage MEDICC2’s tree construction to identify each cell’s most recent common ancestor (Figure 1B). Finally, we infer foreground CNAs by taking bin-wise state differences for each allele between the cell and its MRCA (ForegroundState_bin,allele_ = TotalState_bin,allele_ - MRCAState_bin,allele_) (Figure 1C). After removing events that are ≤ 4 Mb, we use this delta – which represents novel events not reflected by the cell’s ancestry – to assess how drugs impact the CNA profiles of single cells. The pipeline can be retrieved on github and is containerized with singularity (https://github.com/molonc/MRCA_Foreground).

### Feature selection

#### Exploration

We first considered five feature categories comprising 16 total features. Foreground CNA topology was classified as telomere-bounded (spanning from a point in the chromosome arm to the telomere, padded by 2.5 Mb), centromere-bounded (spanning to the centromere, padded by 2.5 Mb), whole-arm (spanning padded centromere to padded telomere), or interstitial (reaching neither boundary). The remaining categories were CNA size: (0.5, 4], (4, 8], (8, 17], (17, 38] and >38 Mb, direction of CN change (gain or loss), magnitude of CN change (1, 2, or 3+), and chromosome type (sex or autosomal). Note that ≤4 Mb events were included here but dropped from later analyses.

We represented each cell by its counts of foreground events across the 16 features, yielding a cell-by-feature matrix. During exploration, we included 17,749 cells across six experimental groups: *TP53*^−/−^ hTERT (2,042 treated and 1,330 untreated), WT hTERT (1,376 treated and 749 untreated), *TP53*^−/−^ BRCA1^−/−^ hTERT (3,473 treated and 3,414 untreated), WT HEK-293 (1,561 treated and 1,188 untreated), SA609 (587 treated and 892 untreated), and SA535 (489 treated and 648 untreated).

Random-forest classifiers were trained separately for each cell line or PDX group containing treated and untreated cells. Within each group, cells were divided into stratified training and held-out test sets at an 80:20 ratio, and missing values were imputed using per-feature medians calculated from the training set. Models comprised 120 trees, with class weights recalculated within each bootstrap sample to account for class imbalance. Trees used Gini impurity, had no maximum depth, required at least two samples to split an internal node and one sample per leaf, and considered the square root of the total number of features at each split.

Performance was evaluated on the held-out test set using receiver operating characteristic area under the curve (ROC–AUC), average precision, accuracy and F1 score. Feature importance was assessed using impurity-based importance and SHAP values calculated for up to 200 held-out test cells per group (Extended Data Figure 2). Features were ranked by mean absolute SHAP value, and signed SHAP values indicated whether higher feature values shifted predictions towards the treated or untreated class. Together, this gave an initial indication that framing the foreground as a feature space can capture drug-induced differences.

#### Foreground Feature space

Motivated by the random forest exploration, and keeping biological interpretability in mind, we settled on defining a foreground feature space (Figure 1D) with: topology (telomere-bounded, centromere-bounded, interstitial, or whole-arm), size: (4, 8], (8, 17], (17, 38] and >38 Mb, and direction of CN change relative to the MRCA (gain or loss). To enforce mutual exclusivity across the feature space, we crossed these categories, treating each combination as a distinct feature (e.g., 4-8 Mb + gain + telomere-bounded is one column in the feature space). This combination spans 32 theoretical categories; 2 small whole-arm combinations do not occur in the human reference genome and are excluded, leaving 30 valid categories. Representing each cell by its event counts across these 30 categories gives the cell-by-category matrix used for decomposition (Figure 1E). Moving forward, we classified foreground CNAs in this feature space and applied HDP signature decomposition across the full dataset.

### Mutational Signature Methods

#### HDP signature decomposition

The foreground of each cell is represented by enumerating the number of foreground CNAs of each feature class as defined by the 30 mutually exclusive features. We removed cells with less than 3 foreground CNA events, leaving us with 43,516 cells. To decompose the foreground space of these cells, we used an HDP implementation called mSigHdp^31^ with the parameters as described in Supplementary Table 7. We started from an initial K of 10, and ran 30,000 burn-in iterations with 6 random restarts. All the cells coming from one sample were grouped under one parent node, and all parent nodes were grouped under one global node (Figure 1F). We used gamma.alpha=1 and gamma.beta=100 for the shape and rate (or scale) hyperparameters for the Gamma distribution prior, which models the concentration parameters of the underlying HDP. Signatures that appeared in fewer than 90% of the Gibbs posterior samples were considered low-confidence and omitted. Signatures that were more similar than cosine similarity 0.9 were merged. We ran 5 seeds with 20 chains each and picked one seed (200437) which was consistent with most other seeds. 2 other seeds produced the same number of signatures (8) with almost identical feature loadings (cosine similarity ≥ 0.99, Supplementary Figure 4B). Two other seeds produced 10 signatures, 8 of which are also found in the picked seed (cosine similarity ≥ 0.99), and 2 other signatures did not contain any additional information. We extracted 200 posterior Gibbs samples (with 100 skipped samples between every one extracted sample and the next, to minimize correlation between extracted samples), and repeated the process for all the 20 chains. All the 200×20 extracted samples were combined together to obtain mutation-to-signature assignments. Of all the input foreground mutational events that characterise a cell (a cell could have, for example, 3 to 100 or more events), HDP assigns a number of these events (the “exposure”) to a specific signature, and may assign other events to another signature.

#### HDP Certainty Assessment

We adapted the methodology established by Nicola Roberts for identifying high-confidence exposures and signatures^77^. Briefly, to determine the confidence of a cell-to-signature assignment, we calculated the lower and upper bounds of the 95% Highest Posterior Density (HPD) credible interval across all posterior MCMC samples from the 20 chains of the selected seed. If the lower bound of this interval is zero, the exposure is deemed not-credible, indicating a lack of confidence that the signature is truly present within that specific sample’s mutational spectra. In this case, we lower the percentage of the HPD credible interval density by 5%, and re-assess if the lower bound is greater than 0. If an assignment passes this criteria, it is said to pass that certainty threshold. While the original hdp implementation (https://github.com/nicolaroberts/hdp) provides built-in functionality for these calculations, the hdpx architecture (https://github.com/steverozen/hdpx) and by extension mSigHdp (https://github.com/steverozen/mSigHdp) does not as it uses a different method for consensus signature derivation. We developed a custom extraction function to perform the HDP interval calculations using the hdpx posterior MCMC samples.

#### PCA of Feature Loadings

We followed standard PCA protocols to assess which features drove separation during signature decomposition.

#### NMF on foreground features

For comparison, we ran a non-negative matrix factorization (NMF) analysis using SigProfilerExtractor^78^ on the same set of samples and features we trained HDP on. We tested a range of total signatures from 1–10, setting the number of NMF replicates to 200, and following manual recommendations set the range of iterations to reach convergence from 10000 to 100000, with the average solution stability set to 0.8, and the minimum solution stability to 0.2. The optimal solution was defined using SigProfilerExtractor’s “estimate best solution”, with default settings.

### Statistical analyses

#### Identifying Whole Genome Duplicated Cells

To identify whole genome duplicated cells, we followed a protocol described by McPherson et al^30^. Briefly, if ≥ 50% of the genome has two or more copies of the major allele, the cell is considered to have undergone at least one WGD event, and with three or more copies a cell is considered to have experienced at least two WGD events.

#### GLMMs to assess differences in mutational burden

To test for differences in event rate counts between drug treatment groups, we fit generalized linear mixed models with negative binomial families. Samples were treated as random effects. Effect sizes were reported as fold changes. P values were corrected with Bonferroni within drug comparisons. We used the same model to assess differences in the mutational burden of the feature space. When less than 3 samples were available, we took cell-wise means for evaluating signature differences and reported number of cells and number of samples.

#### GLMMs to assess differences in signature exposure

To quantify differences in mutational signature activity between treatment groups, per-cell signature exposures were derived from HDP decomposition. Cellular exposure to signatures were weighted by their certainty (cellular exposure x certainty), such that signatures with low certainties were down-weighted. For each drug comparison, for each signature, we fit a univariate generalized linear mixed model with a Tweedie family. Samples were treated as random effects, and contrasts of interest were extracted from the model. The p values of the model’s coefficients were corrected with Bonferroni across all signatures for each drug comparison. When less than 3 samples were available, we took cell-wise means for evaluating signature differences and reported number of cells and number of samples.

#### Cell to signature assignment for proportional comparisons

To quantify differences in how many cells were affected by each signature across different samples, we assigned cells to signatures by assigning the cell to whichever signature (certainty threshold ≥ 0.5) it was most highly exposed to. If no signature passed the certainty threshold, it was not assigned. As previously mentioned, cells that were not assigned signatures were included in the proportional calculations. This includes cells that had less than 3 foreground CNAs (with notable inclusion of cells that had zero foreground events), cells identified as S-phase, and cells with no signature assignment that passed the certainty threshold.

## Data Availability

All data are available for general research use. Processed data, including topology-annotated allele-specific copy number profiles for DLP+, and foreground feature counts per cell (input matrix for HDP) is available at https://doi.org/10.5281/zenodo.22016654. Raw scWGS BAMs for cell lines and newly published CX5461 treated SA535 are available for download at https://ega-archive.org/studies/EGAS50000002035.

## Code Availability

The pipeline which wraps SIGNALs and MEDICC2, extracts foreground, and annotates CNA topology is available here: https://github.com/molonc/MRCA_Foreground. Code for HDP, statistical tests, and figure generation are available here: https://doi.org/10.5281/zenodo.22016654. DLP+ processing pipeline is available here: https://github.com/molonc/mondrian_nf, https://github.com/bpotvin-bccrc/mondrian_utils (forked from https://github.com/mondrian-scwgs/mondrian).

## Acknowledgements

We want to acknowledge the contribution of Daniel Lai, who led the project for a substantial amount of time prior to moving on to a new job.

## Author Contributions

S.A. conceived and supervised the study and wrote the manuscript. M.A. led the bioinformatic analysis and developed the HDP signature model. A.A.-H. performed bioinformatic analyses, contributed to pipeline development and produced the main figures. D.Y. performed the cell line experiments. W.D. developed the foreground extraction model. D.L. led bioinformatic processing and developed the foreground calling algorithm. D.Y., A.A.-H., M.A. and S.A. contributed biological interpretation and wrote the manuscript.

B.F. implemented WGD calling and NMF signature extraction. E.Z. performed genotype validation and background copy number analyses. A.S. performed feature importance and PERT analyses. S.M. helped develop the exposure uncertainty code. J.C. wrote the feature extraction code. J.T. performed variant calling analyses. H.T. performed preliminary transcriptomic analyses.

D.Y., H.X., B.M., N.W. and J.Z. performed cell line and validation experiments; C.O., V.C. and B.K. generated drug sensitivity data; F.K. and T.R.D.A. performed xenograft experiments. V.A., A.R., M.V.V., C.B. and B.W. prepared DLP+ libraries and sequenced cells; R.R., J.M., B.P.-P., and S.B. ran the Mondrian pipelines and managed data. R.M. and A.M. supported sequencing. All authors reviewed and approved the manuscript.

---

**Extended Data Figure 1.**
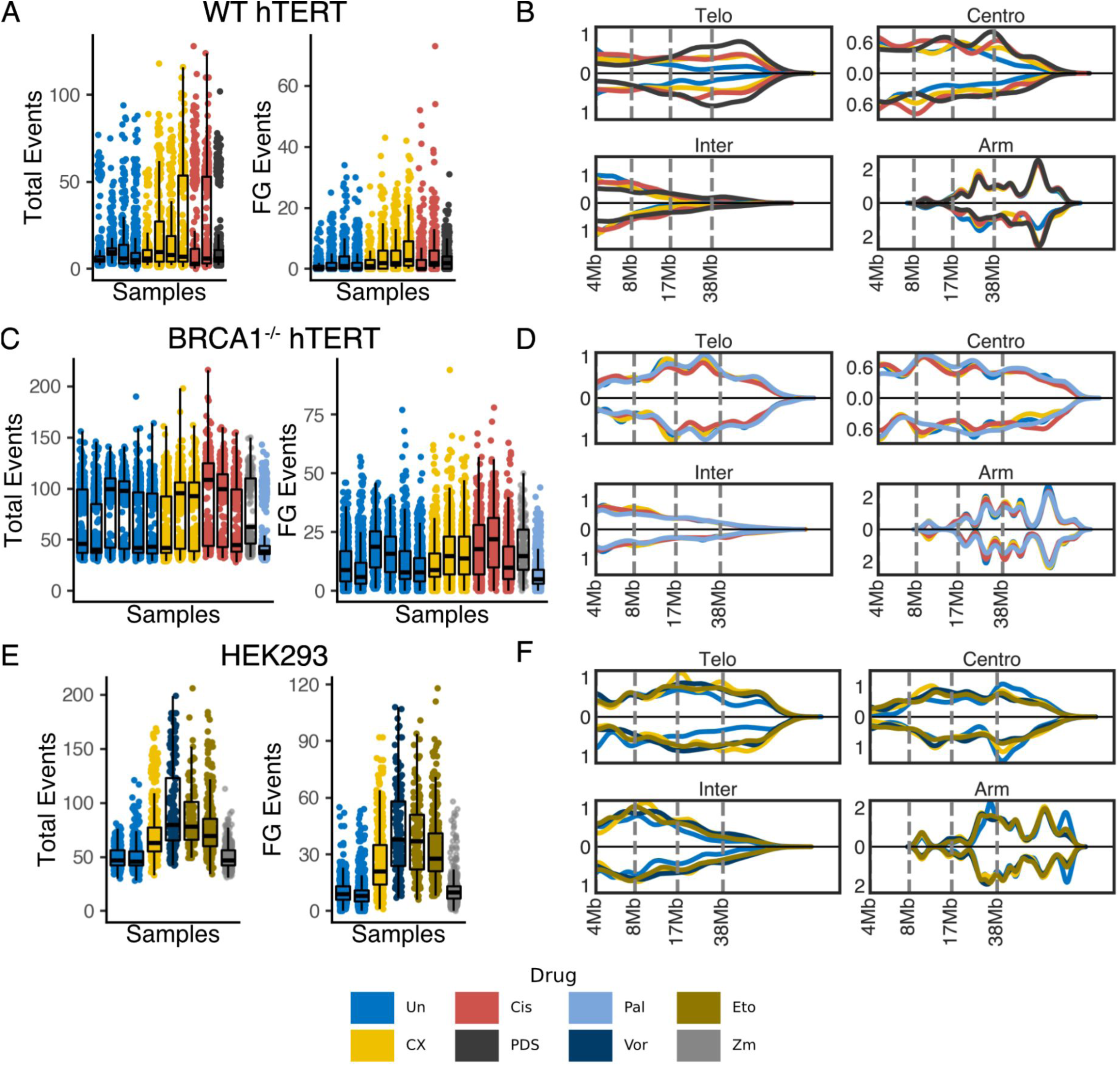
Event Rate and Per-Cell Feature Density of other cell lines and drug treatments. **A,** Total and foreground event rates per cell across WT hTERT samples from a variety of drug treatments. **B,** CNAs are categorized into 30 mutually exclusive features. Density of feature space counts per cell for WT hTERTs. **C, D,** A-B for *TP53⁻/⁻BRCA1⁻/⁻*-deficient hTERTs. **E, F,** A-B for HEK-293 cells.

**Extended Data Figure 2.**
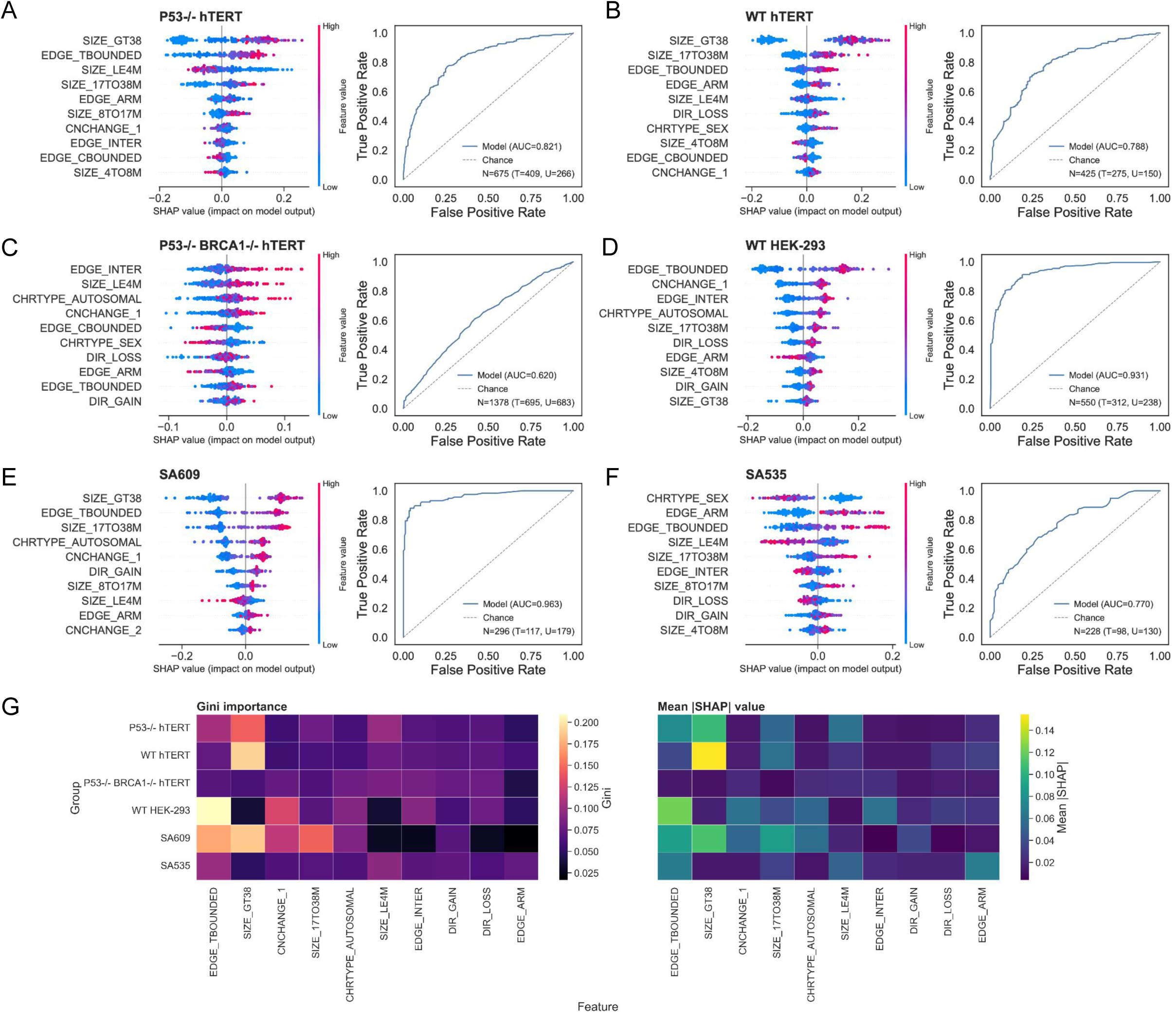
Random forest analysis of uncrossed feature importance. **A–F,** SHAP beeswarm plots and ROC curves for random forest models distinguishing treated from untreated cells in *TP53*^−/−^ hTERT (**A**), WT hTERT (**B**), *TP53^−/−^BRCA1*^−/−^ hTERT (**C**), WT HEK-293 (**D**), SA609 (**E**) and SA535 (**F**). SHAP plots show the ten most important uncrossed features in each group. Each point represents one cell; colour indicates the feature value, and horizontal position indicates its contribution to the model prediction. Positive SHAP values favour classification as treated. ROC curves show classification performance, with the dashed diagonal indicating chance and the AUC reported in each panel. (**G),** Heatmaps comparing feature importance across groups using Gini importance (left) and mean absolute SHAP value (right). Greater colour intensity indicates higher feature importance.

**Extended Data Figure 3.**
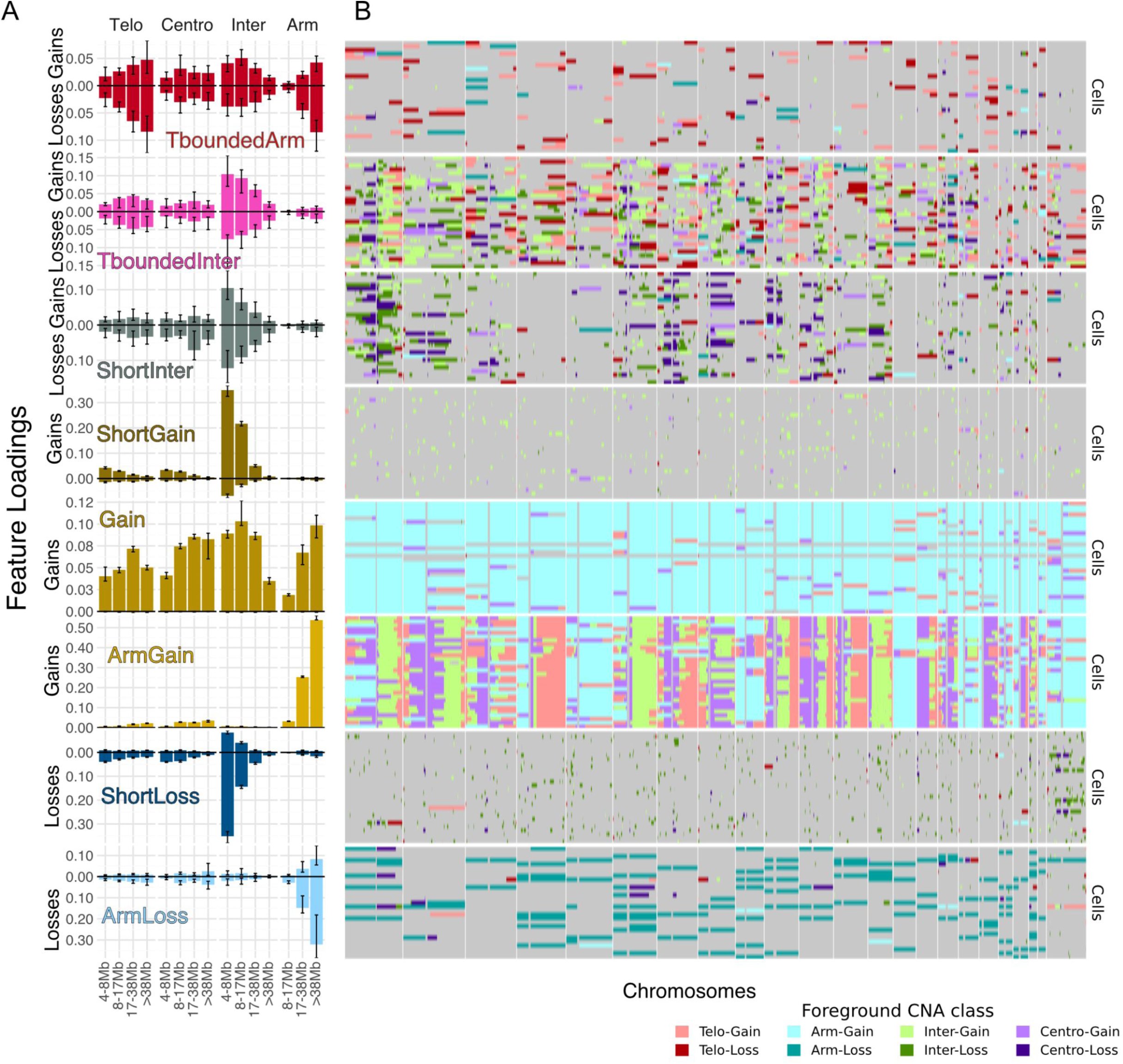
Foreground Signature Loadings and Top Examples. **A,** Loadings of our 30 features on the 8 signatures discovered by HDP trained on 43,516 single cell whole genomes. Signatures were named after their highly loaded features. Loadings on gain-related features are above the x axis, whereas loadings on loss-related features are below. Error bars represent 95% credible intervals. **B,** Foreground CNA profiles of the top 20 cells loaded with each signature in A. Gains and losses are denoted by lighter and darker colors respectively. Red colors represent telomerically-bounded CNAs, blue - whole arm, green - interstitial, and purple - centromerically-bounded.

**Extended Data Figure 4.**
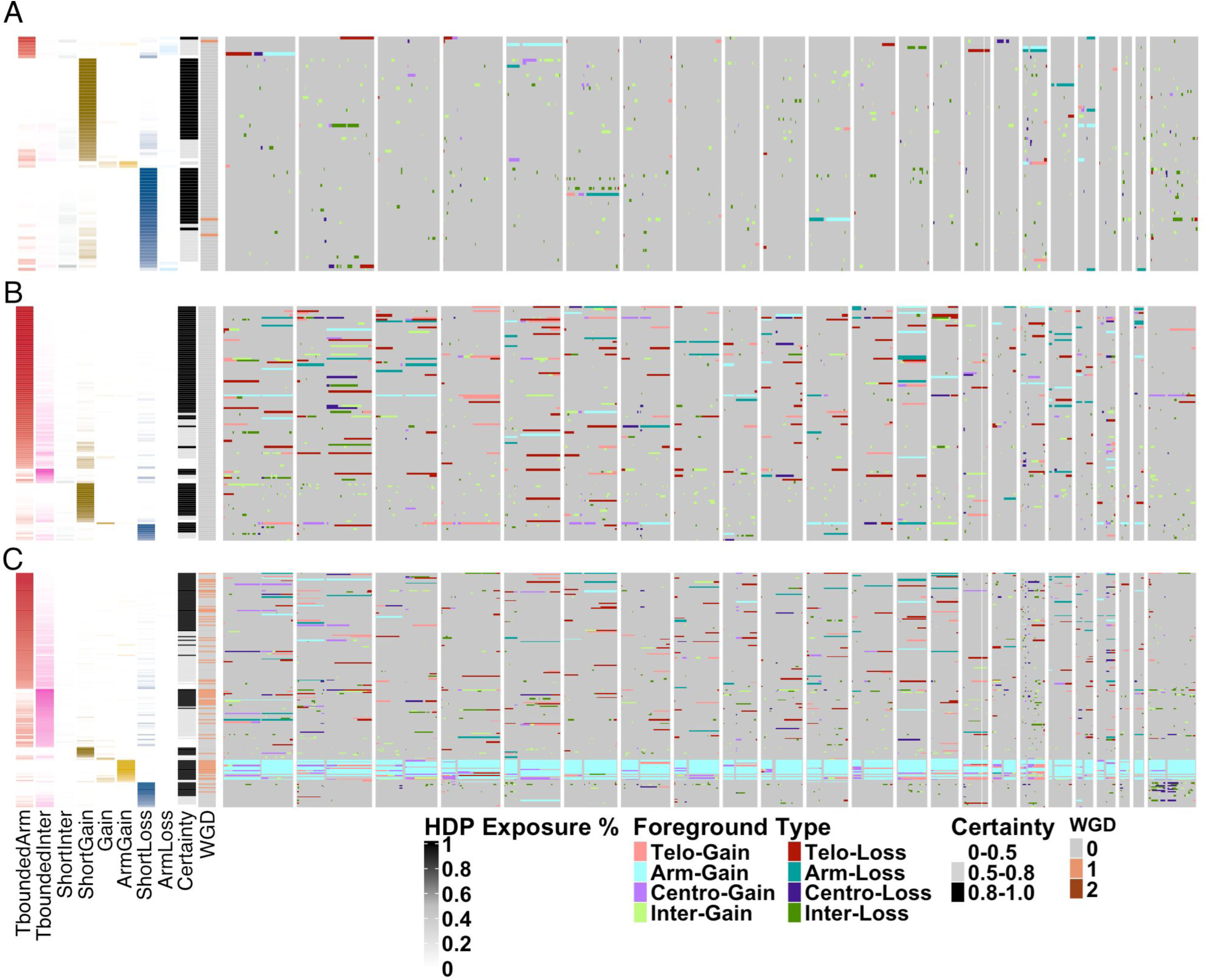
Foreground heatmap for selected TP53^-/-^ hTERT cells. **A**, untreated. **B**, treated with CX5461 for 7 days at 703nM. **C**, treated with cisplatin for 7 days at 1144nM. Each row shows the combined foreground from the major and minor alleles of a cell, annotated by signature, certainty and WGD. Columns are positions in the genome, organized by chromosomes. Color coded are foreground CN segments, classified into categories depending on the topology and the foreground CN direction.

**Extended Data Figure 5.**
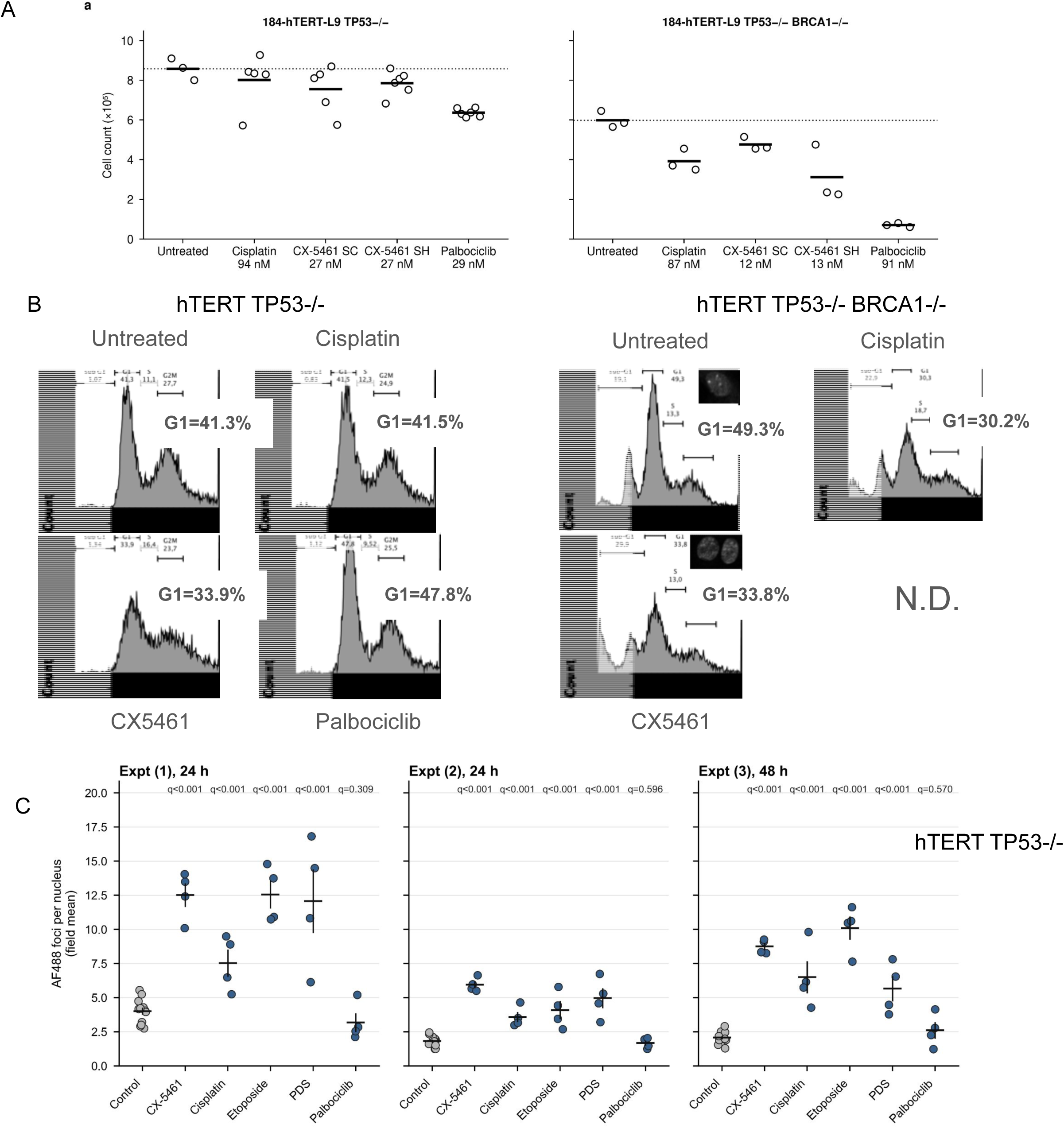
Cell number, Cell Cycle and 53BP1 foci assessment in treated hTERT cells. Treatment of hTERT *TP53^-^*^/-^ (left) and hTERT *TP53^-/-^BRCA1*^-/-^ (right) cells with the indicated concentration of drugs and **A.** cell count of replica wells after 7 days and **B.** representative cell cycle profiles after 7 day treatment in the conditions as in (A). **C.** Treatment of hTERT *TP53^-/^*^-^ with 700 nM CX5461, 1.2 µM Cisplatin, 1 µM Etoposide or 6.24 µM PDS results in DNA damage after 24-48 hr as assessed by 53BP1 (AlexaFluor-488nm, AF488) nuclear foci. Treatment with solvent or 50 nM Palbociclib (CDK4/6 inhibitor) does not induce DNA damage foci as assessed by the same method. Chart shows the mean number of 53BP1 spots per nuclei (n=4 fields counted) for the labelled 5 treatments across 3 experiments, and q-values shown are Benjamini-Hochberg false-discovery-rate-adjusted across 15 planned treatment-versus-control comparisons.

**Extended Data Figure 6.**
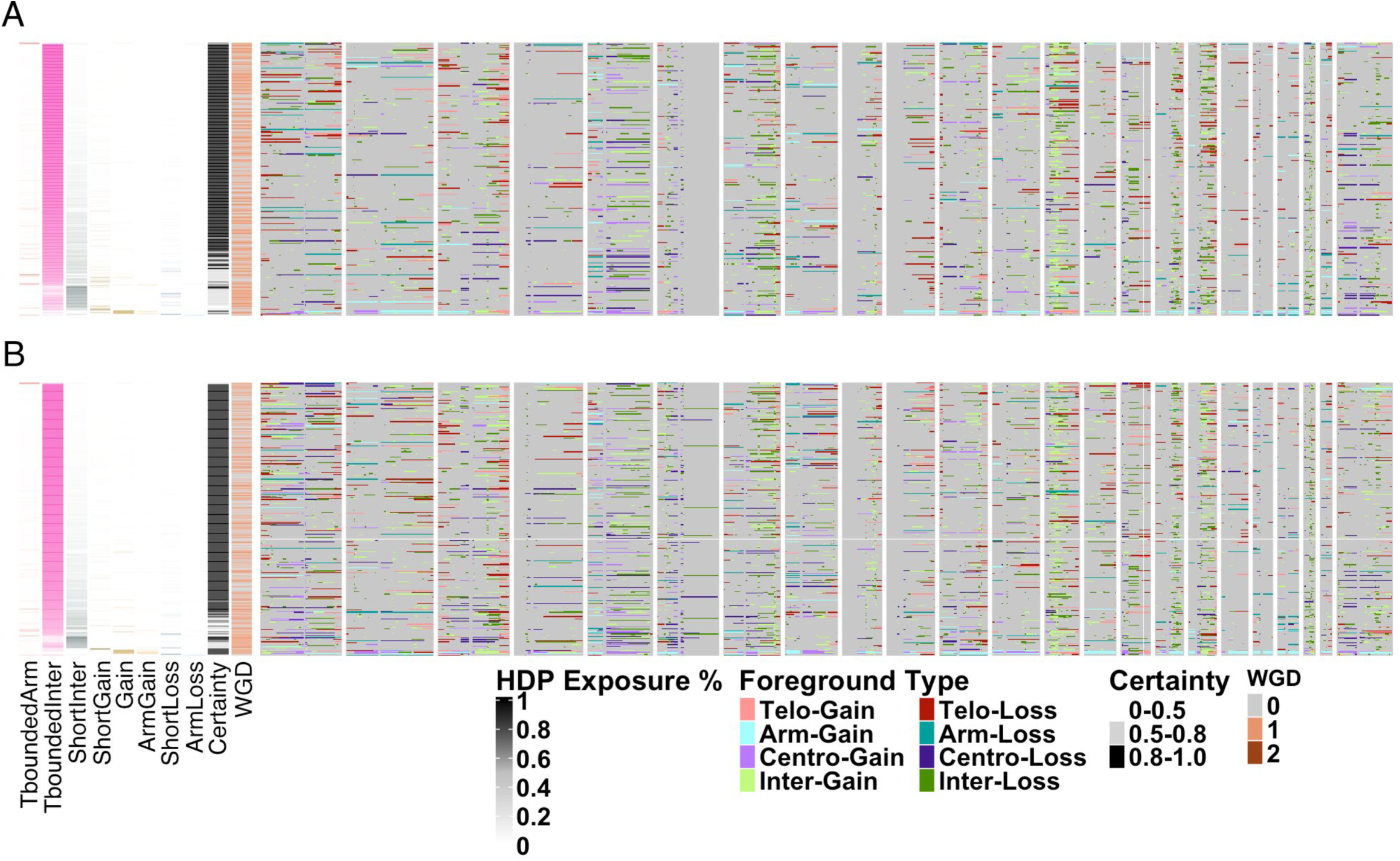
Foreground heatmap for selected TP53^-/-^ BRCA1^-/-^ hTERT cells. **A**, untreated. **B**, treated with CX5461. Each row shows the combined foreground from the major and minor alleles of a cell, annotated by signature, certainty and WGD. Columns are positions in the genome, organized by chromosomes. Color coded are foreground CN segments, classified into categories depending on the topology and the foreground CN direction.

**Extended Data Figure 7.**
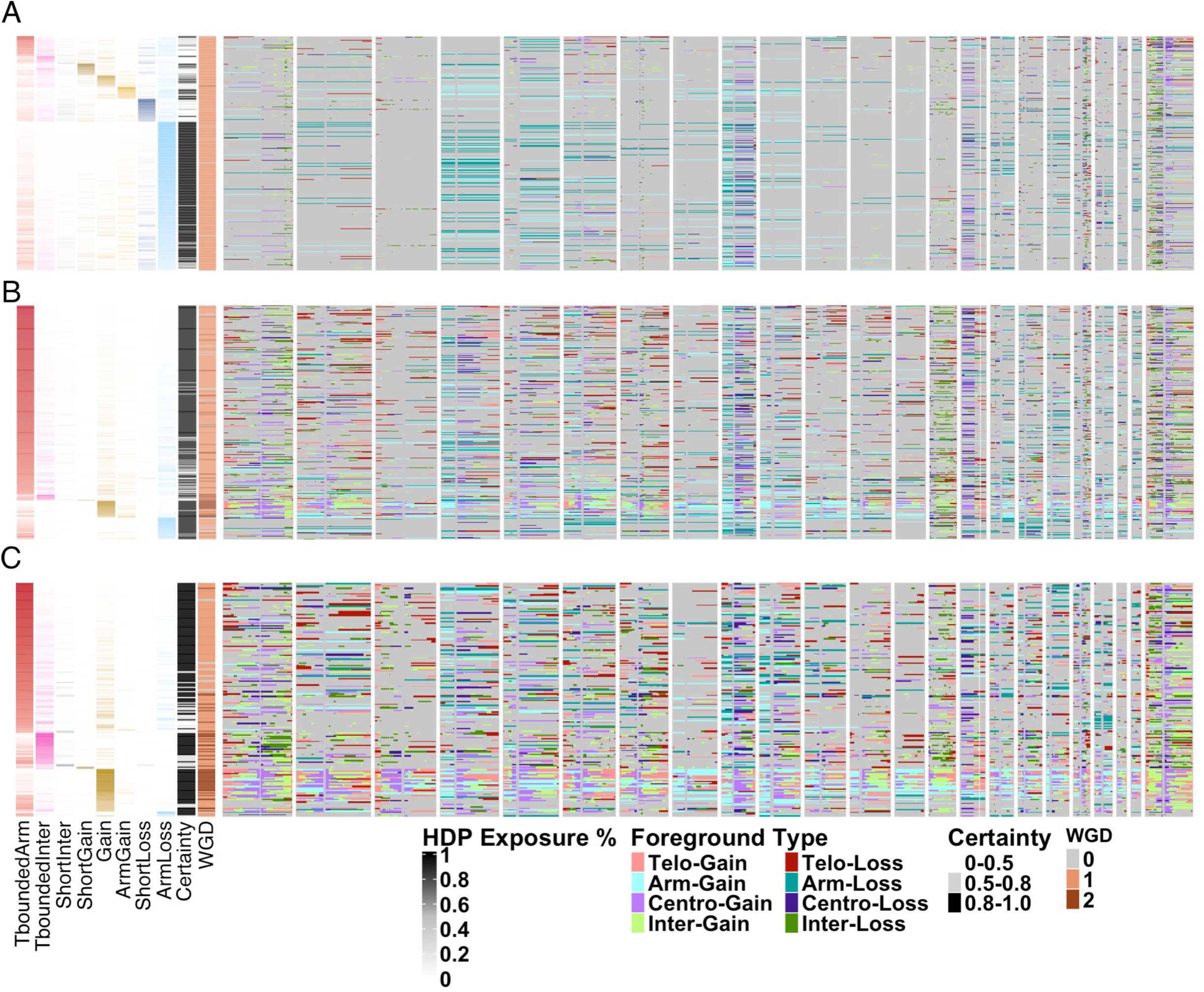
Foreground heatmap for selected wild type HEK-293 cells. **A**, untreated. **B**, treated with CX5461. **C,** treated with etoposide. Each row shows the combined foreground from the major and minor alleles of a cell, annotated by signature, certainty and WGD. Columns are positions in the genome, organized by chromosomes. Color coded are foreground CN segments, classified into categories depending on the topology and the foreground CN direction.

**Extended Data Figure 8.**
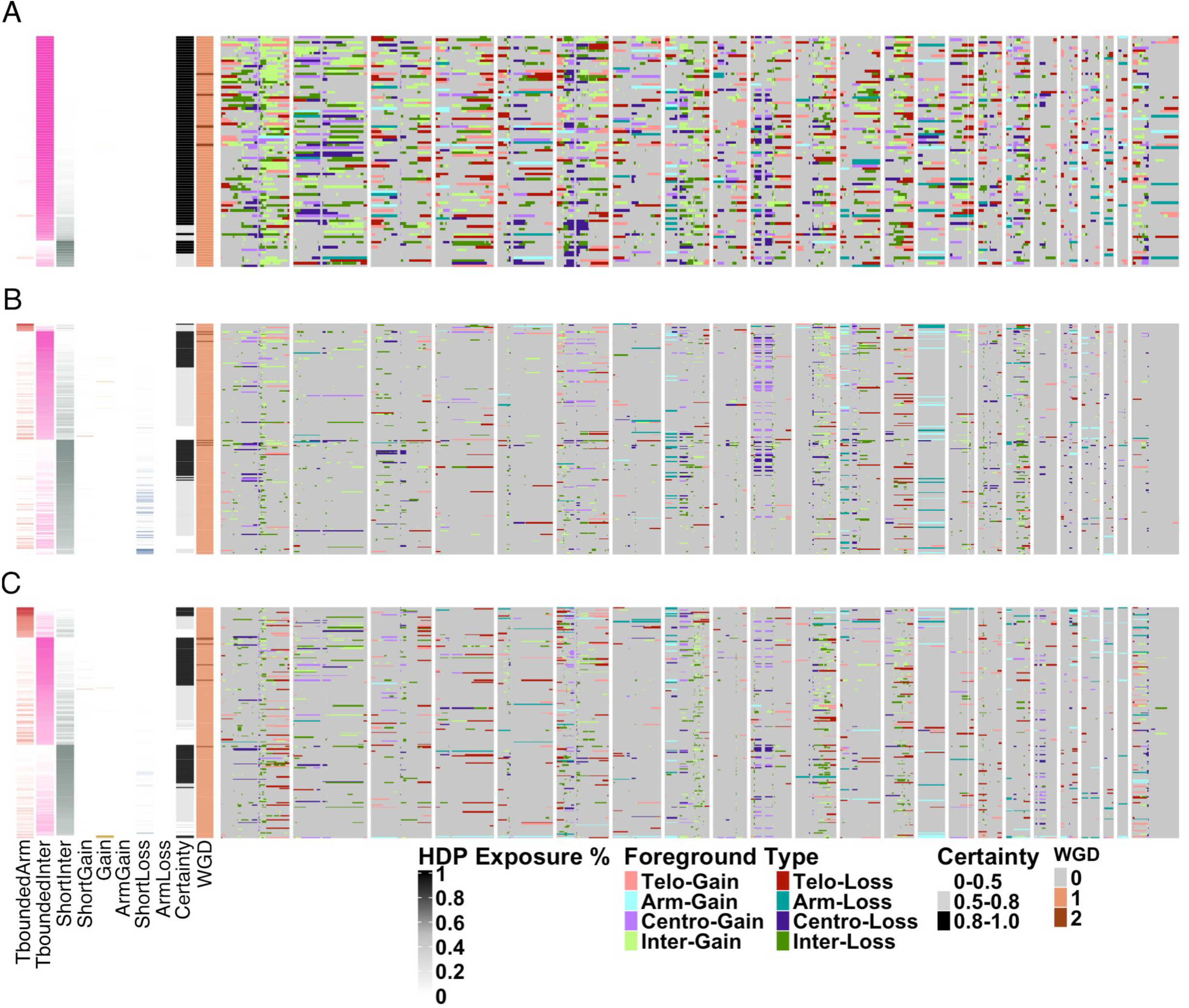
Foreground heatmap for selected PDX SA609. **A**, untreated at generation 1. **B**, treated with cisplatin for 1 generation. **C,** treated with cisplatin for 1 generation followed by 1 untreated generation. Each row shows the combined foreground from the major and minor alleles of a cell, annotated by signature, certainty and WGD. Columns are positions in the genome, organized by chromosomes. Color coded are foreground CN segments, classified into categories depending on the topology and the foreground CN direction.

**Extended Data Figure 9.**
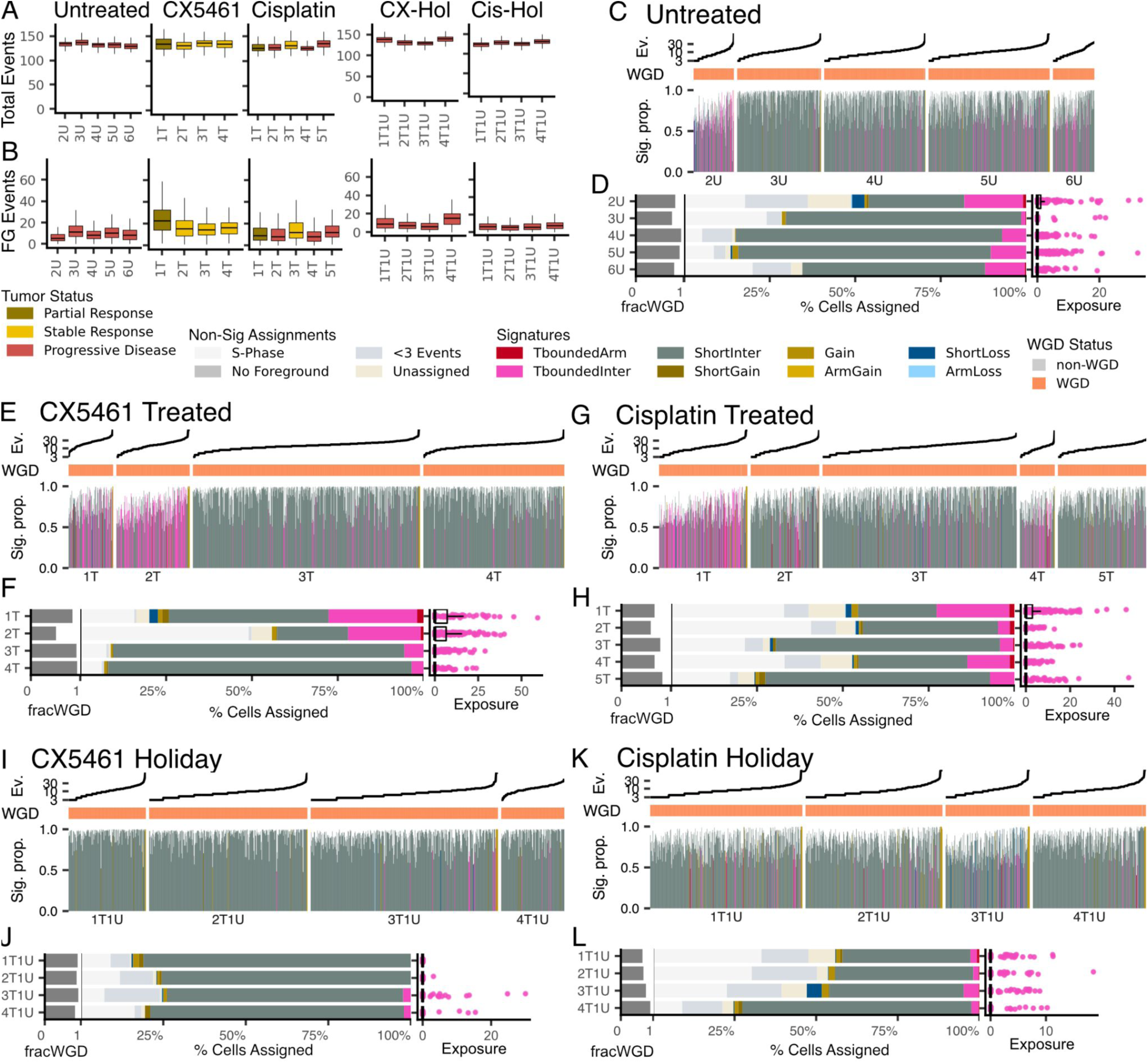
Foreground mutational signatures in SA535 PDX branches. SA535, a breast carcinoma PDX line, was propagated along five branches: treated with cisplatin or CX5461 at each transplant generation, left untreated across all generations, or treated with cisplatin or CX5461 and then given one drug-holiday generation. Generations are labeled with a number followed by T (treated) or U (untreated); for example, 1T is the first, treated generation. **A,** Event rates of absolute mutational CNAs for the branches. Tumor status indicates response to drug at the time of sequencing. **B,** Foreground event rates. **C,** Per-cell proportional exposure to each signature in the untreated branch, with each column representing one cell. Events with <0.5 certainty in assignment are included in the percentage calculation but plotted as white space. Cell-wise event rate and whole-genome doubling (WGD) status are annotated above the plot. **D,** Proportion of cells assigned to each signature, or the reason they were filtered out prior to signature decomposition. The left margin shows the fraction of WGD cells per library; the right margin shows cell-wise exposure to TboundedInter. **E,F,** Exposures for the CX5461-treated branch**. G,H,** Exposures for the cisplatin-treated branch. **I,J,** Exposures for the CX5461-holiday branch. **K,L,** Exposures for the cisplatin-holiday branch.

**Supplementary Figure 1.**
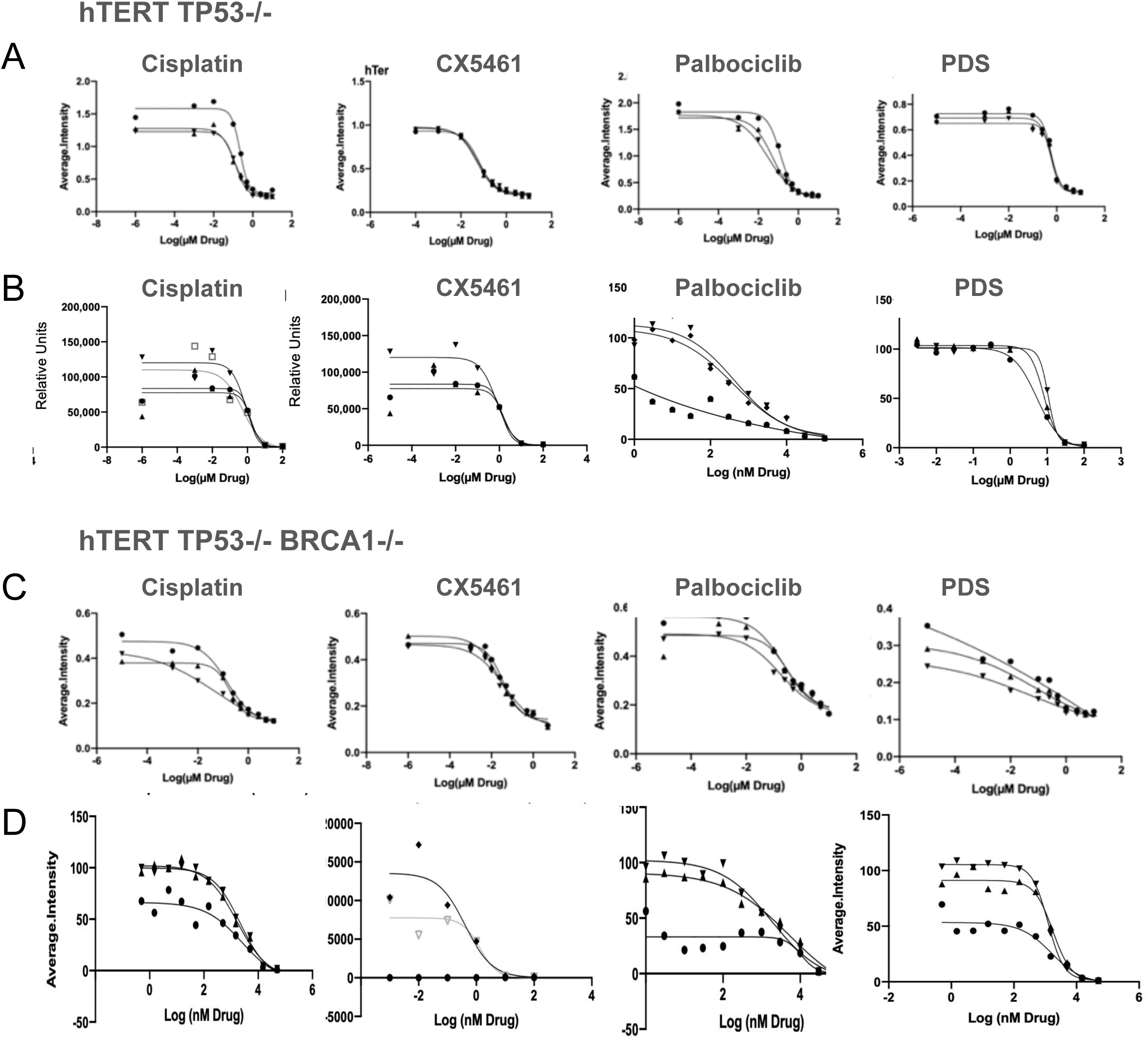
IC50 for hTERT TP53^−/−^ and TP53-/-BRCA1-/- cell lines. Representative dose–response curves from independent experiments used to determine IC50, which is defined as the concentration producing 50% growth inhibition (50% viability) relative to untreated controls, derived from 4-parameter logistic fits under the assay and time point indicated, for each compound using 5-day Colony forming assay, (CFA stained with crystal violet, CV) for **A.** TP53-/- and **C.** TP53-/-BRCA1-/-. or 3 or 5-day CellTiter-Glo (CTG) for **B.** TP53-/- and **D.** TP53-/-BRCA1-/-. Points represent biological replicates (n=3 or 4).

**Supplementary Figure 2.**
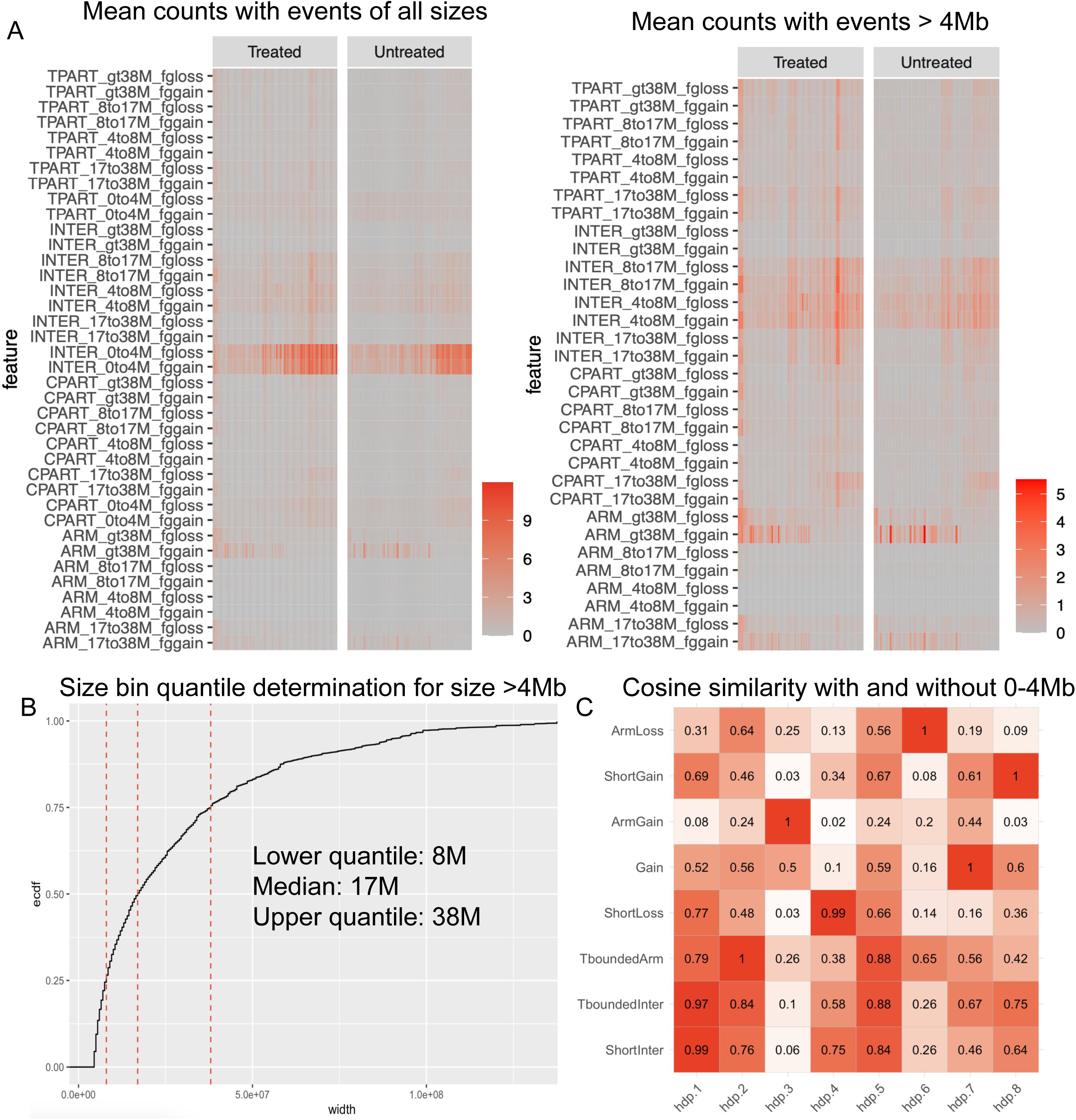
Analysis of the impact of the size feature. **A** left, mean counts for treated and untreated samples for 38 crossed features: 4 topology bins (TPART = telomere bounded, INTER = interstitial, CPART = centromere-bounded, ARM = whole arm), 5 size bins, and 2 foreground direction bins (gain/loss). The counts are dominated by the INTER_0to4M features. Right, mean counts, but after removing all the features with size 0-4Mb, where the feature counts are more equally distributed. **B**, after removing events 4Mb and less, the bin sizes are determined by taking the lower and upper quantiles of the data. **C**, cosine similarity between the signatures in this manuscript on the y axis (Figure 2F) and the signatures obtained by also including the 0-4Mb bin size on the x axis.

**Supplementary Figure 3.**
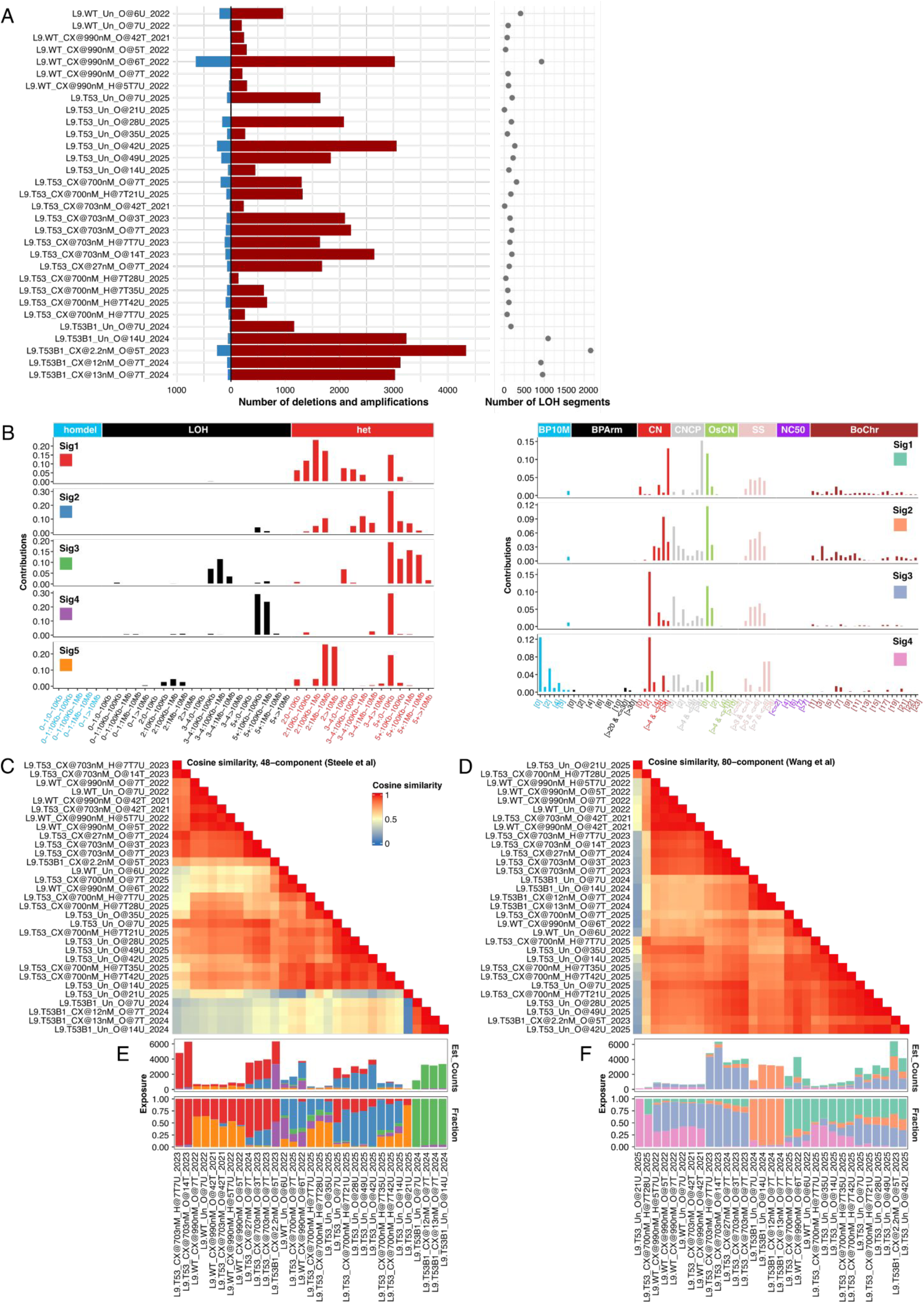
Bulk CNA distribution in hTERT cells. **A,** The total number of deletion and amplification segments (left) and the absolute number of LOH segments (right) across pseudobulked hTERT cells. **B**, Composition of CNA signatures extracted from the pseudobulk data using Steele et al features and Wang et al features. Cosine similarity of samples based on the distribution of Steele et al (**C**) and Wang et al (**D**) CNA features. Absolute and relative exposures of the NMF-extracted CNA signatures for Steele et al (**E**) and Wang et al (**F**) classification approaches.

**Supplementary Figure 4.**
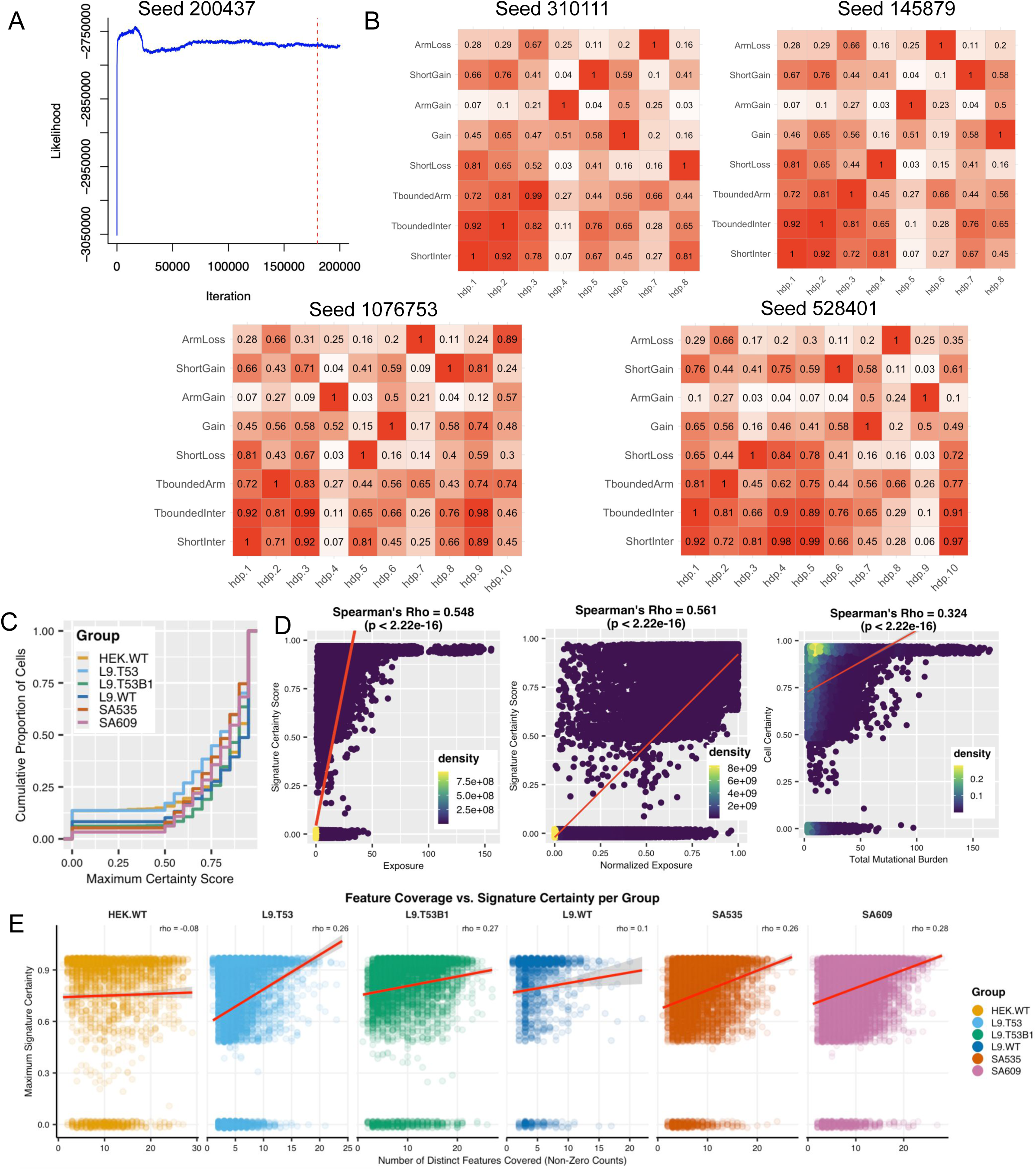
HDP stability and certainty analysis. **A,** Likelihood plot for one HDP chain (total number of chains run = 20) for seed 200437, which was the seed selected in this manuscript (Figure 2F). For each chain, 180,000 iterations were run for burn-in (30,000 iterations x 6 random resets), up to the dashed vertical red line, followed by posterior sampling, where the likelihood remained stable. **B**, cosine similarity between the selected seed and 4 other seeds. The signatures in the selected seed (y axes) were found with cosine similarity ≥ 0.99 in other seeds (x axes). **C**, cumulative proportion of cells in each group, by certainty score. **D**, signature/cell certainty versus exposure, normalized exposure or total mutational burden. **E**, Signature certainty versus the number of distinct features

**Supplementary Figure 5.**
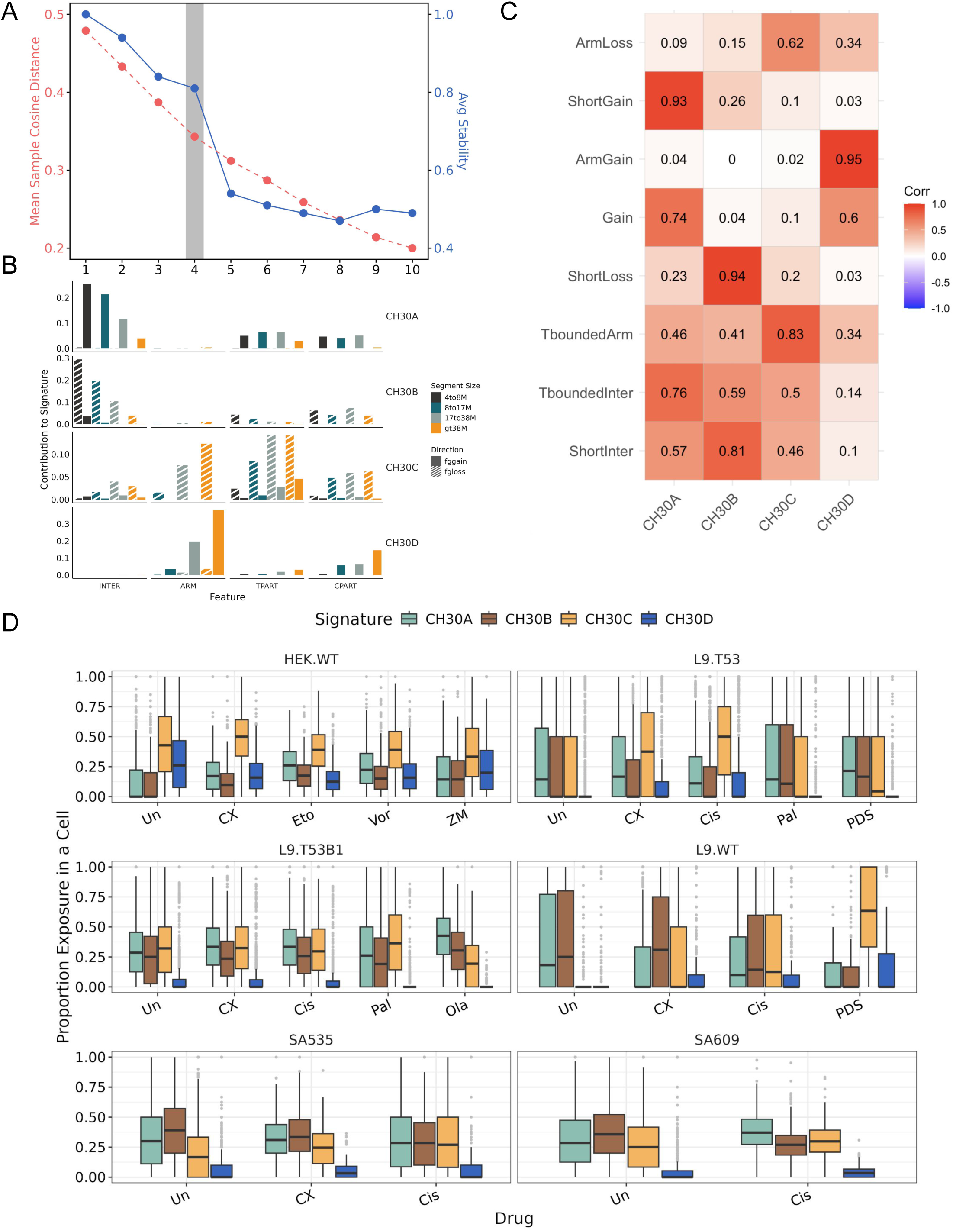
Non-negative matrix factorization (NMF) analysis of foreground CN features. **A**, Total signature selection done by SigProfilerExtractor. **B**, The four optimal signatures found by NMF analysis. **C**, Cosine similarity to the eight optimal HDP signatures. **D**, Cell-wise signature exposure proportions across assessed cell lines and experimental conditions. Drug names on x-axis: CX - CX5461, Eto - etoposide, Vor - voreloxin, PDS - pyridostatin, Ola - Olaparib, Cis - cisplatin, Pal - palbociclib, Un - untreated.

**Supplementary Figure 6.**
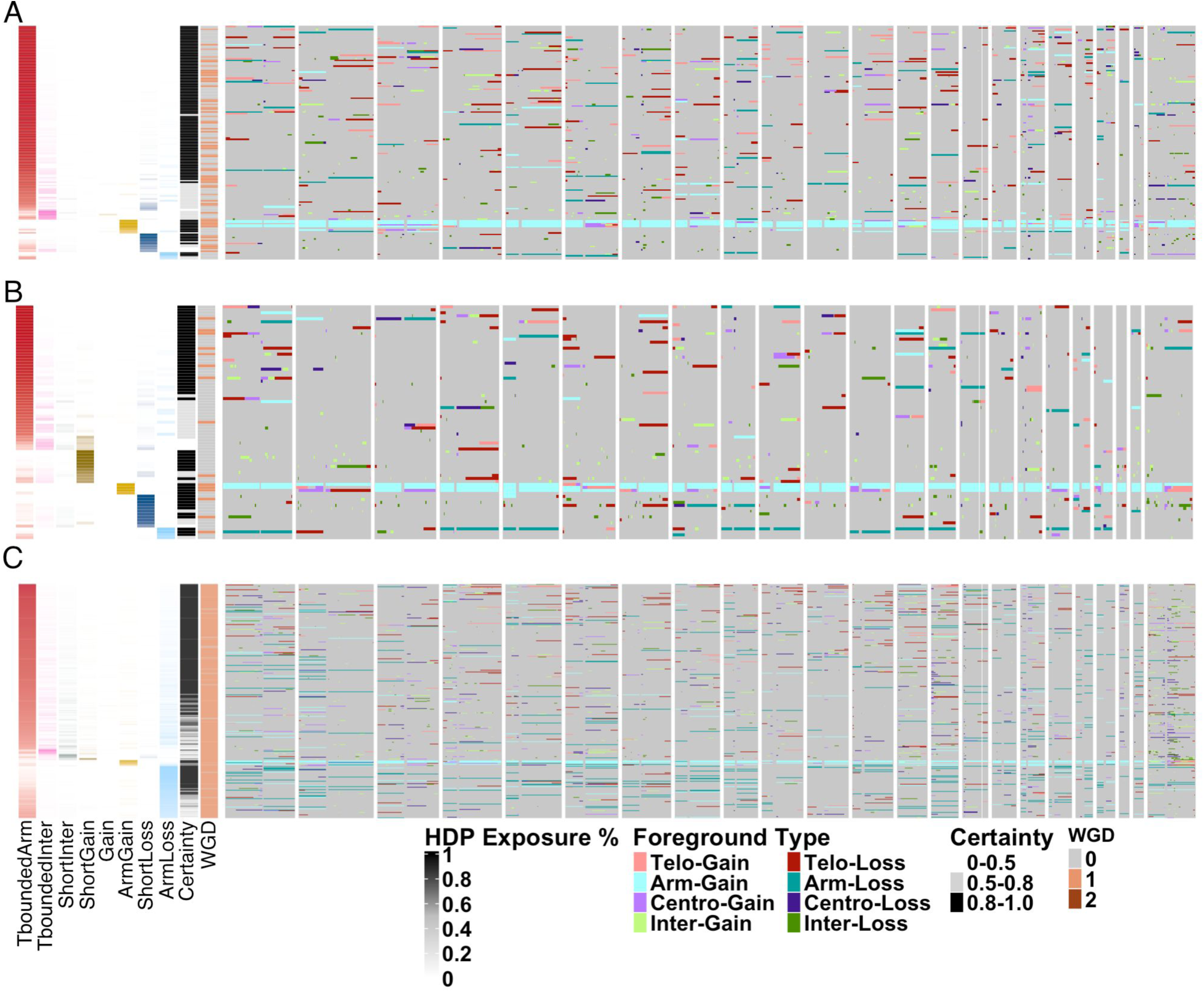
Samples from TP53^-/-^ hTERT cisplatin drug persistence line. **A**, treated with cisplatin for 7 days. **B**, treated for 7 days, followed by 7-day holiday. **C**, treated for 7 days, followed by 35-day holiday. Each row shows the combined foreground from the major and minor alleles of a cell, annotated by signature, certainty and WGD. Columns are positions in the genome, organized by chromosomes. Color coded are foreground CN segments, classified into categories depending on the topology and the foreground CN direction.

**Supplementary Figure 7.**
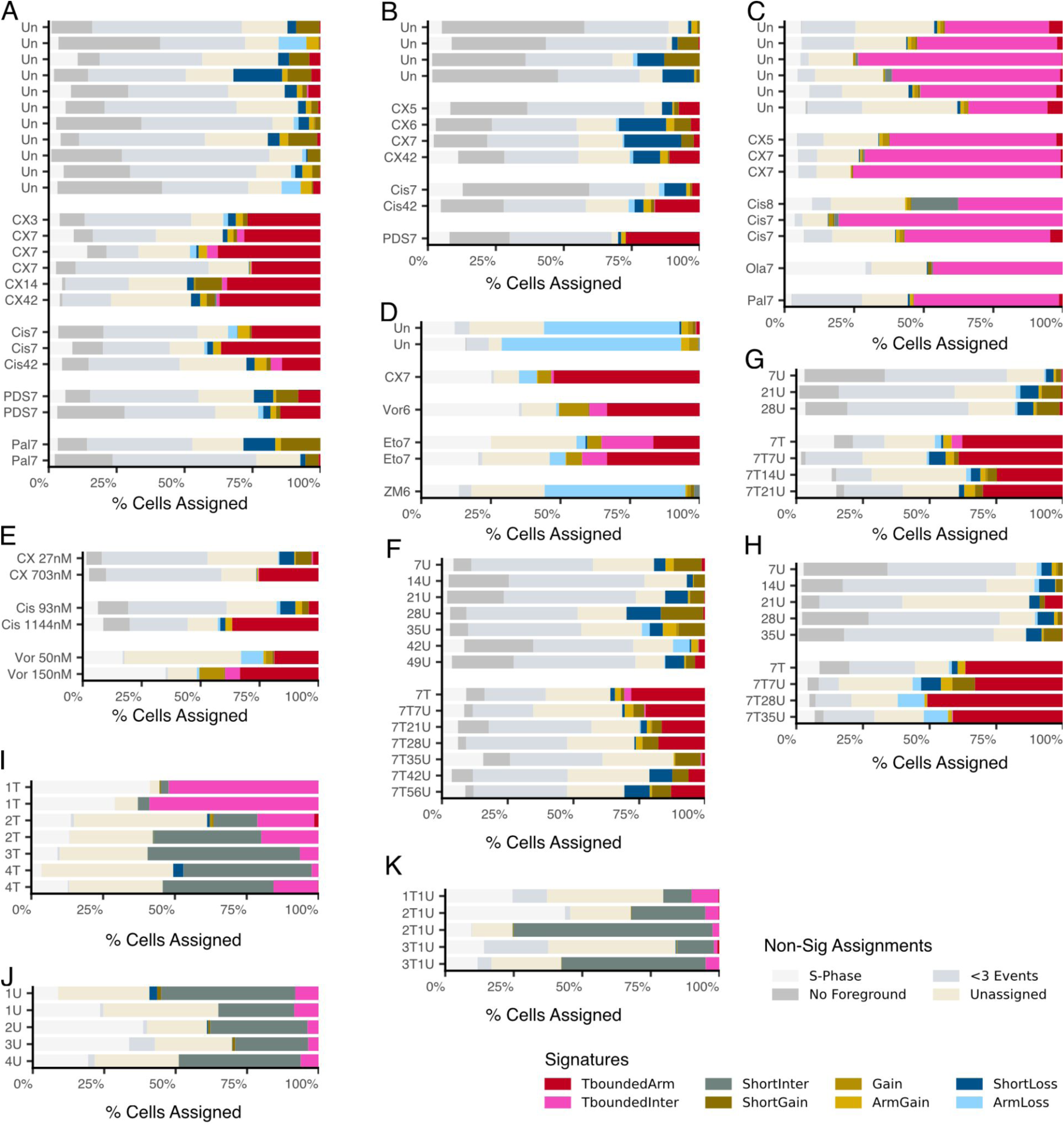
Cell-Signature Assignments at Certainty Threshold of 0.8. Proportion of cells assigned to each signature at 0.8 certainty threshold, or the reason they were filtered out prior to signature decomposition. **A,** TP53-deficient hTERTs. **B,** WT hTERTs. **C,** BRCA1-deficient hTERTs. **D,** WT HEK293s. **E,** Dose-dependent comparisons. **F,** Line 1 of the TP53-deficient hTERTs CX5461 holiday study. **G,** Line 2 of the TP53-deficient hTERTs CX5461 holiday study. **H,** TP53-deficient hTERTs cisplatin holiday study. **I,** Cisplatin-treated SA609. **J,** Untreated SA609. **K,** Holiday SA609.

**Supplementary Table 1 - IC50 of drugs on cell lines.**

- Drug: Name of small molecule assayed
- Endpoint: Reporting endpoint - Inferred valued based on graph: Hill Slope, IC50. IC30 is calculated.
- Rep N: Means from each biological replicate experiment
- Mean: The means from all experiments
- SD: Standard Deviation from all experiments
- Unit: Concentration of the drug
- Mean (nM): Standardised concentration of the drug in nM
- Dose used (unit): Unit and dose of the drug used in the experiments.
- Experimental Conditions: Type and length of assay to measure cell viability

**Supplementary Table 2 - GLMM coefficients for absolute and foreground event rate.**

- genotype: Genotype of cell line
- DRUG_NAME_SHORT: Drug treatment of model comparison
- n_treatment: Number of treated libraries
- n_reference: Number of untreated libraries
- Coef: GLMM coefficient for drug-treatment effect
- SE: Standard Error of coefficient
- CI_2.5: Lower confidence bound of Coef
- CI_97.5: Upper confidence bound of Coef
- Fold change: mean fold-increase of drug-treated cells compared to untreated, derived from Coef
- P_value: p value of coef
- P_adj: Bonferroni adjusted p-value. P’s adjusted within each genotype
- Significant: True significant, alpha = 0.05

**Supplementary Table 3 - Samples included in the random forest classification.**

- DESC_NAME: a unique identifier that describes the sample
- CELL_LINE_ORIGIN: cell line name or patient identifier for PDX
- GROUP: a short identifier that groups samples from the same cell line, genotype and/or patient
- DRUG_NAME: drug name or untreated for no treatment
- DRUG_REGIME: number of treatments days (T), number of holiday days (U) where applicable, or NA for no treatment
- DRUG_DOSE: dose used for treatment, or NA for no treatment
- METRICS_CELLS: number of sequenced cells in total
- HQ_CELLS: number of cells with quality score ≥ 0.75 and S-phase probability < 0.5
- N_SPHASE: total number of S-phase cells independent of quality measure
- HQ_SPHASE: number of S-phase cells with quality measure ≥ 0.75
- MEDICC_INPUT: number of cells that were used as input to medicc2
- CELLS_W_FG: number of cells with at least one foreground event
- NCELLS_MIN3: number of cells with at least three foreground events

**Supplementary Table 4 - Random forest SHAP values.**

- Genotype: Genotype or cell line random forest is trained on
- EDGE_INTER: SHAP values from random forest for separating treated and untreated cells, for feature characterizing if foreground CNA has an interstitial topology
- EDGE_TBOUNDED: Same as previous, for feature characterizing if foreground CNA has a telomere-bounded topology
- EDGE_CBOUNDED: Same as previous, for feature characterizing if foreground CNA has a centromer-bounded topology
- EDGE_ARM: Same as previous, for feature characterizing if foreground CNA has a whole arm topology
- SIZE_LE4M: Same as previous, for feature characterizing if foreground CNA is less than 4Mb in size
- SIZE_4TO8M: Same as previous, for feature characterizing if foreground CNA is 4MB to 8Mb in size
- SIZE_8TO17M: Same as previous, for feature characterizing if foreground CNA is 8MB to 17Mb in size
- SIZE_17TO38M: Same as previous, for feature characterizing if foreground CNA is 17MB to 38Mb in size
- SIZE_GT38: Same as previous, for feature characterizing if foreground CNA is greater than 38Mb in size
- DIR_GAIN: Same as previous, for feature characterizing if foreground CNA is a gain
- DIR_LOSS: Same as previous, for feature characterizing if foreground CNA is a loss
- CNCHANGE_1: Same as previous, for feature characterizing if foreground CNA is change of 1 relative to MRCA
- CNCHANGE_2: Same as previous, for feature characterizing if foreground CNA is change of 2 relative to MRCA
- CNCHANGE_3PLUS: Same as previous, for feature characterizing if foreground CNA is change of 3 relative to MRCA
- CHRTYPE_SEX: Same as previous, for feature characterizing if foreground CNA is on sex chromosome
- CHRTYPE_AUTOSOMAL: SHAP values from random forest for separating treated and untreated cells for feature characterizing if foreground CNA is on autosomal chromosome

**Supplementary Table 5 - GLMM coefficients for signature exposures.**

- Same as Supplementary Table 2

**Supplementary Table 6 - Samples used in HDP.**

- DESC_NAME: a unique identifier that describes the sample
- CELL_LINE_ORIGIN: cell line name or patient identifier for PDX
- GROUP: a short identifier that groups samples from the same cell line, genotype and/or patient
- DRUG_NAME: drug name or untreated for no treatment
- DRUG_REGIME: number of treatments days (T), number of holiday days (U) where applicable, or NA for no treatment
- DRUG_DOSE: dose used for treatment, or NA for no treatment
- METRICS_CELLS: number of sequenced cells in total
- HQ_CELLS: number of cells with quality score ≥ 0.75 and S-phase probability < 0.5
- N_SPHASE: total number of S-phase cells independent of quality measure
- HQ_SPHASE: number of S-phase cells with quality measure ≥ 0.75
- MEDICC_INPUT: number of cells that were used as input to medicc2
- CELLS_W_FG: number of cells with at least one foreground event
- NCELLS_MIN3: number of cells with at least three foreground events
- NCELLS_ANALYSED: number of cells with that have at least three foreground events of size > 4M bases

**Supplementary Table 7 - HDP hyperparameters.**

- Parameter: hyperparameters used for HDP training
- Value: value of hyper parameter

**Supplementary Table 8 - HDP Signatures.**

- Feature: Feature name
- ShortInter: Loadings of Feature for the ShortInter signature
- TboundedInter: Loadings of Feature for the TboundedInter signature
- Rest of columns: Loadings of Feature for the given signature

**Supplementary Table 9 - PDX Genotype annotations.**

- PDX sample: PDX sample ID
- Pathogenic somatic variant alleles: “X”, calculated by “Y”
- Germline variants:
- CHORD probability HRD
- CHORD probability BRCA1
- Funnell classification:

